# Changes in an Enzyme Ensemble During Catalysis Observed by High Resolution XFEL Crystallography

**DOI:** 10.1101/2023.08.15.553460

**Authors:** Nathan Smith, Medhanjali Dasgupta, David C. Wych, Cole Dolamore, Raymond G. Sierra, Stella Lisova, Darya Marchany-Rivera, Aina E. Cohen, Sébastien Boutet, Mark S. Hunter, Christopher Kupitz, Frédéric Poitevin, Frank R. Moss, Aaron S. Brewster, Nicholas K. Sauter, Iris D. Young, Alexander M. Wolff, Virendra K. Tiwari, Nivesh Kumar, David B. Berkowitz, Ryan G. Hadt, Michael C. Thompson, Alec H. Follmer, Michael E. Wall, Mark A. Wilson

## Abstract

Enzymes populate ensembles of structures with intrinsically different catalytic proficiencies that are difficult to experimentally characterize. We use time-resolved mix-and-inject serial crystallography (MISC) at an X-ray free electron laser (XFEL) to observe catalysis in a designed mutant (G150T) isocyanide hydratase (ICH) enzyme that enhances sampling of important minor conformations. The active site exists in a mixture of conformations and formation of the thioimidate catalytic intermediate selects for catalytically competent substates. A prior proposal for active site cysteine charge-coupled conformational changes in ICH is validated by determining structures of the enzyme over a range of pH values. A combination of large molecular dynamics simulations of the enzyme in crystallo and time-resolved electron density maps shows that ionization of the general acid Asp17 during catalysis causes additional conformational changes that propagate across the dimer interface, connecting the two active sites. These ionization-linked changes in the ICH conformational ensemble permit water to enter the active site in a location that is poised for intermediate hydrolysis. ICH exhibits a tight coupling between ionization of active site residues and catalysis-activated protein motions, exemplifying a mechanism of electrostatic control of enzyme dynamics.

## Introduction

The role of enzyme dynamics in catalysis is one of the most intensively investigated topics in modern biophysics ^1, 2, 3^. Enzymes, like all proteins, exist in conformational ensembles corresponding to multiple populated minima in their potential energy landscapes ^4, 5^. Protein dynamics are a combination of motions within these minima and transitions among them. The conformations that compose an ensemble can have intrinsically different catalytic proficiencies, permitting functional selection of specific conformations during catalysis ^6, 7^, in the laboratory ^8, 9^, or through evolution ^10, 11, 12^. Because the catalytic cycle transiently changes the underlying protein energy landscape, enzyme conformational ensembles also change during catalysis ^7, 13^. Although equilibrium or near-equilibrium enzyme ensembles are more accessible to study using biophysical approaches, understanding how enzymes facilitate their remarkable rate enhancements ultimately requires directly observing their non-equilibrium behavior during catalysis.

Time-resolved X-ray crystallography can be used to study the non-equilibrium behavior of proteins. Pioneering work using Laue crystallography ^14, 15, 16, 17, 18^ paved the way for a new generation of time-resolved experiments that use serial X-ray crystallography at X-ray free electron laser (XFEL) ^19, 20^ or synchrotron ^21, 22^ sources to follow the responses of proteins to perturbation. By collecting data from thousands of microcrystals, serial crystallography permits crystal manipulation by soaking in substrates ^23, 24^, activation by light ^25, 26^, temperature jump ^27^, or releasing caged substrates ^28, 29^ as well as dramatically reducing radiation damage effects ^30, 31^. Application of time-resolved crystallography and computational methods to several enzymes has revealed that catalysis can activate enzyme motions not apparent in the absence of substrate ^32, 33^. However, interpretation of the difference electron density maps produced by time-resolved crystallography experiments can be challenging ^34^. Minor conformations populated stochastically rather than in concert during catalysis are especially difficult to identify and model. Because these minor conformations can play significant roles in catalysis ^35^ and in contributing to protein entropy ^36^, it is imperative to develop strategies to enrich them to study important regions of the reaction coordinate.

Our enzyme of interest is isocyanide hydratase (ICH), which catalyzes the irreversible hydration of isocyanides to N-formamides via formation of a cysteine-derived thioimidate intermediate (Fig. 1A) ^32, 37, 38, 39^. Isocyanides (also called isonitriles) possess a triple bonded nitrogen-carbon (R-N^+^≡C^-^) functional group in resonance with a double-bonded electrophilic carbenoid species (R-N=C). These compounds often possess antibiotic and chalkophore (copper-binding) activities ^40, 41, 42, 43, 44^. ICHs are commonly found in pseudomonad bacteria and soil-dwelling fungi and likely serve a defensive role against isocyanide natural products produced by competing microbes ^38, 45^. In addition to its intrinsic biochemical interest, ICH is a new model system for catalysis-activated enzyme dynamics. Prior XFEL mix-and-inject serial crystallography (MISC) studies of *Pseudomonas* ICH showed that formation of the thioimidate intermediate causes the enzyme to populate a different conformational ensemble during catalysis ^32^. In wild-type ICH, covalent modification of the active site cysteine thiolate led to distributed changes in the ICH conformational ensemble across the dimer, especially for helix H (residues 152-166) near the active site in protomer A ^32^ (Fig. S1). However, the isomorphous (F_o_-F_o_) difference electron density around mobile regions of the enzyme was difficult to interpret on its own, and structures of ICH with an oxidized Cys101 nucleophile that enriched these conformations were needed to confirm a structural model where helix H moved upon intermediate formation owing to a loss of negative charge on the Cys101 thiolate. Two mutations (G150T and G150A) that reduced ICH catalytic rate approximately 6-fold were also engineered for that study. The 277 K crystal structure of G150T ICH showed that helix H is shifted, similar to a minor conformation sampled by wild-type ICH when Cys101 is modified by formation of the catalytic intermediate ^32^. Because G150T stably adopts a state that is transiently populated by the wild-type enzyme, it is a useful tool for studying how ICH conformational ensembles change during the catalytic cycle.

**Figure 1:**
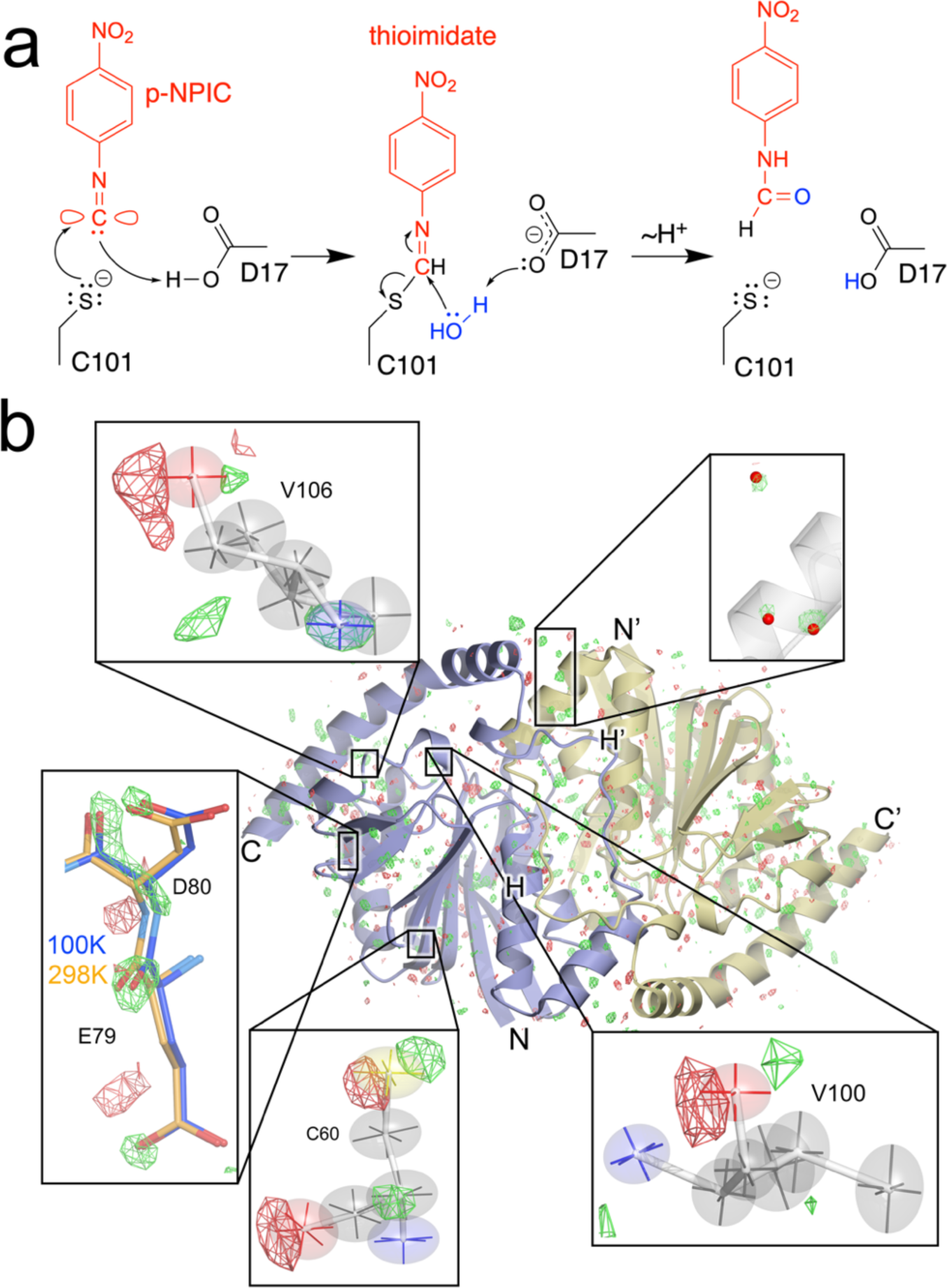
ICH mechanism and evidence for temperature-dependent modes of intrinsic enzyme flexibility. (a) Postulated ICH mechanism is shown for the para-nitrophenylisocyanide (p-NPIC) substrate used in this study. The isocyanide is shown in its carbenoid form, consistent with the electrophilic character needed for attack by the catalytic Cys101 thiolate nucleophile in ICH. Formation of the thioimidate intermediate is postulated to be facilitated by the general acid Asp17, which protonates the C1 atom and then later acts as a general base to activate water (blue) for thioimidate hydrolysis. (b) Temperature-dependent changes in the ICH dimer. The F_o_(274K synchrotron)-F_o_(298K XFEL) isomorphous difference electron density map is contoured at 3 σ, with positive features shown in green and negative features in red. Insets show regions of special interest, including the alignment of difference electron density features along the principal axes of anisotropic ADPs in Cys60, Val100, and Val106. F_o_(274K synchrotron)-F_o_(298K XFEL) peaks near ordered waters (red spheres, top) indicate higher occupancy of these waters at 274K than 298K. For residues 79 and 80, the difference electron density indicates displacements that agree with the structural differences observed in a 100 K crystal structure (blue bonds) compared to the 298 K XFEL structure (gold bonds).

Here we use mix-and-inject serial crystallography (MISC)^23, 24^ at the Linac Coherent Light Source (LCLS) XFEL to observe how catalysis affects the conformational ensemble of G150T ICH. The high resolution (1.3 Å) of the G150T ICH XFEL MISC data permits building detailed structural models for the resting enzyme and the catalytic intermediate as well as refinement of anisotropic atomic displacement parameters (ADPs) for both structures. Changes in the protonation of active site residues during catalysis modulates G150T ICH conformational dynamics and this mutant provides a clearer view of a key step in ICH catalysis than was possible with the wild-type enzyme. The relevance of these results to time-resolved studies of other enzymes is discussed.

## Results

### XFEL and synchrotron crystal structures of G150T ICH have differences that lie along intrinsic modes of enzyme flexibility

We obtained a 1.3 Å resolution XFEL structure of resting G150T ICH at 298 K using 20-30 μm microcrystals delivered via a MESH injector at the MFX beamline at LCLS (see Methods). The refined XFEL structure of G150T ICH superimposes nearly exactly (0.14 Å all-atom RMSD) with prior 1.1-1.2 Å resolution 274 K synchrotron structures of this mutant protein ^46^, as expected. However, there are surprising differences between the XFEL and synchrotron structures evident in F_o_(synchrotron)-F_o_(XFEL) isomorphous difference electron density maps, which contain over 300 peaks at 3.2σ (0.2 e^-^/Å^3^) or higher (Fig. 1b). By comparison, a F_o_-F_o_ map of two replicate synchrotron G150T datasets has no features above 0.1 e^-^/Å^3^, establishing that the F_o_(synchrotron)-F_o_(XFEL) peaks are genuine signal. Several of the largest positive difference peaks (5-6σ) are near ordered solvent and indicate higher occupancy of these waters in the 274 K synchrotron dataset (Fig. 1b). Other peaks are near residues sampling multiple conformations (e.g. Cys67 and Cys101) and suggest higher occupancies for minor conformers in the 298 K XFEL dataset (Fig.S2). In cases where positive and negative F_o_(synchrotron)-F_o_(XFEL) peaks are paired near an atom, these features often lie roughly along a principal axis of the anisotropic atomic displacement parameter (ADP) ellipsoid. This was seen most prominently near well-ordered and electron dense atoms, such as peptide oxygen and sulfur atoms in Val100, Val106, and Cys60 (Fig. 1b). The correspondence between these difference map features and the directional preferences of the anisotropic ADPs suggests that small atomic displacements between the synchrotron and XFEL datasets occur along intrinsically preferred directions of atomic motion in the protein.

The movement of atoms along “softer” internal degrees of freedom in the enzyme is further supported by comparison of a cryogenic (100 K) structure with the 274K synchrotron and 298 K XFEL structures of G150T ICH. The cryogenic structure superimposes on the XFEL structure of G150T with all-atom RMSD of 0.447 Å (Cα RMSD=0.205 Å) and many F_o_(synchrotron)-F_o_(XFEL) map peaks lie in the direction of displacements between the cryogenic and XFEL structures (Fig. 1b). We observe these difference peaks in F_o_(synchrotron)-F_o_(XFEL) maps calculated from three independent 274 K G150T ICH synchrotron datasets ^46^, demonstrating reproducibility (Fig. S3). A plausible explanation for these difference electron density features is the 24°C difference in temperature between the synchrotron and XFEL datasets, which can be presumed to shift conformational equilibria. Minor radiation damage to the synchrotron dataset or differences in the sample environment during synchrotron and XFEL data collection cannot be ruled out as potential contributing factors but are unlikely to be dominant effects ^47, 48^. Significant differences between room-temperature and cryogenic crystal structures are well-documented ^49, 50, 51, 52, 53^ and protein conformational ensembles can respond in complex ways as temperature is increased ^54^. Consistent with these prior studies, our results indicate that even moderate changes in temperature that are well above the “glass transition” at ∼180 K leave a clear imprint on protein conformational heterogeneity in atomic resolution isomorphous difference electron density maps. Moreover, our observation that these features often correlate with preferred directions of atomic motion inferred from anisotropic ADPs supports the physical validity of the directional information in the anisotropic ADPs.

### The resting conformational ensemble of G150T ICH is sampled by wild-type ICH during active site protonation and catalysis

The G150T ICH structure overlaps well with features in the isomorphous difference electron density maps calculated using previously collected wild-type ICH XFEL data before and after formation of the thioimidate intermediate (Fig. S4), demonstrating that G150T ICH natively occupies a state that wild-type ICH samples transiently upon catalytic intermediate formation. In prior work, we proposed that the thioimidate intermediate neutralizes the negative charge of Cys101 thiolate, modulating a network of H-bonds that facilitate conformational changes across the ICH dimer ^32^. By contrast, G150T ICH samples this conformational ensemble at rest because of steric effects of the G->T substitution at residue 150 that are distinct from electrostatic changes at the active site in the wild-type enzyme. Because thioimidate formation transiently adds a bulky group to Cys101 during catalysis, it seemed plausible that some previously observed changes in wild-type ICH conformational ensemble might be due to steric effects rather than changes in Cys101 charge. To test the hypothesis that neutralizing the negative charge on the C101 thiolate is sufficient to cause wild-type ICH to sample G150T-like conformations, we determined synchrotron crystal structures at 100 K of wild-type ICH at various pH values in the range of 4.2 to 8.3. We hypothesized that the Cys101 thiolate would become protonated at lower pH values, recapitulating the loss of the negative charge on the Cys101 Sγ atom caused by covalent bond formation during catalysis. Consistent with these predictions, pH values below 6.0 cause a shift in helix H in wild-type ICH that closely resembles the resting G150T ICH structure (Fig. 2a). Wild-type ICH structures at pH 5.4 and 6.0 contain both strained and shifted helix H conformations while structures at pH 5.0 and 4.2 show only the shifted helix conformation, suggesting that pH values in the range of 5-6 are near the Cys101 pK_a_ value (Fig. 2b; Fig. S5, S6). This inference is supported by the pH dependence of ICH activity, which has a maximum at pH 5.0-5.5 (Fig. 2c). This value agrees well with the pK_a_ values of homologous cysteine residues in related DJ-1 superfamily proteins ^55, 56, 57^, suggesting that the Cys101 thiol pK_a_ is in the 5-6 range. Therefore, using pH as a tool to perturb ICH conformational ensembles shows that helical mobility in ICH is closely correlated with Cys101 charge state and that the G150T mutant recapitulates the wild-type ICH conformational ensemble when Cys101 is protonated or covalently modified. Although the G150T mutant, wild-type ICH at low pH, a previously characterized C101A mutant ^37^ and Cys101-SOH oxidized ICH ^32^ all sample similar ensembles with a shifted helix H, only G150T is catalytically active. The activity of G150T ICH is essential for this work because it allows us to characterize changes in its conformational ensemble during catalysis and to relate these to infrequently sampled conformations of the wild-type enzyme.

**Figure 2:**
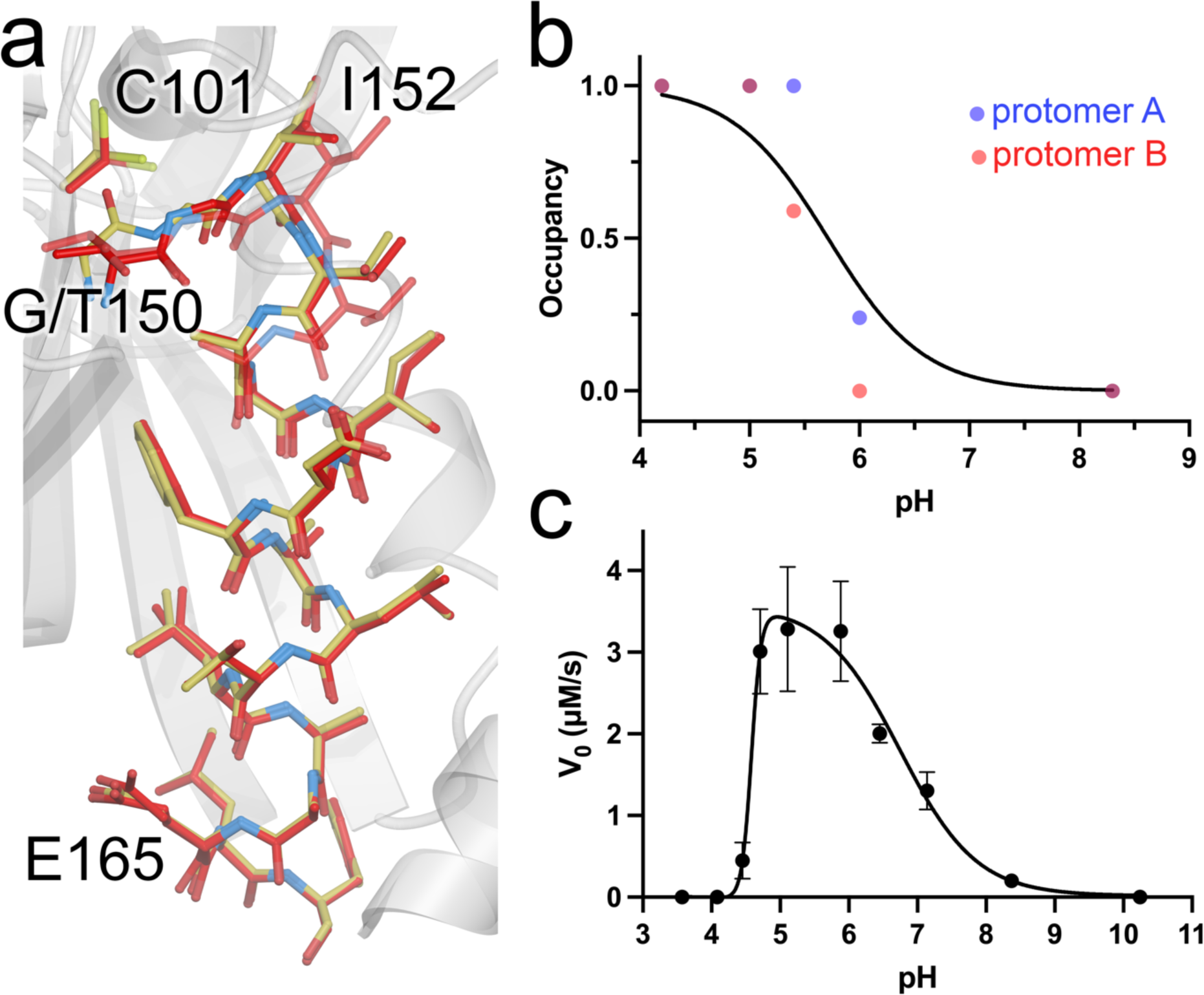
G150T ICH predominantly populates conformations that are sampled by wild-type ICH upon Cys101 charge neutralization. (a) Close structural agreement between the shifted conformation of helix H in resting G150T at pH 8.8 (red) and wild-type ICH at pH 4.2 (gold). Alternate conformations are shown as semi-transparent bonds. (b) Helix H conformational changes owing to Cys101 protonation. The refined occupancies of the shifted (i.e. G150T-like) conformation of helix H in crystal structures of wild-type ICH are plotted against pH. ICH crystallizes with two molecules in the asymmetric unit, with occupancies of helix H for protomer A shown as blue points and protomer B as red. The data are fitted to the Henderson-Hasselbalch equation (black; see Methods) with an apparent pK_a_ of 5.7. (c) pH vs. rate profile for wild-type ICH. Data were measured in triplicate with error bars showing standard error of the mean and fitted using a dose-response curve with a rising inflection point at 4.6, maximum at 5.0, and falling inflection point of 6.7.

### Substrate selects catalytically competent G150T ICH active site conformations

To observe catalysis in G150T ICH microcrystals, we performed a MISC experiment by mixing the substrate para-nitrophenyl isocyanide (p-NPIC) with G150T ICH microcrystals using the coMESH injector and determined a 1.3 Å resolution XFEL crystal structure at 298 K. Like wild-type ICH ^32^, we found that larger (>100 μm) G150T ICH crystals are damaged by the introduction of substrate, necessitating the serial microcrystallography approach taken here. The resting structure of G150T has pronounced conformational disorder at the active site, providing an opportunity to observe how this ensemble of conformations responds to catalysis. In the resting state of G150T ICH, the Cys101 nucleophile populates two major conformations, one corresponding to the conformation observed in wild-type ICH and another that would conflict with Ile152 in the resting wild-type ICH conformation. The shifted helix H in G150T ICH opens a pocket that permits this conformational heterogeneity at Cys101 (Fig. 3a). After 30 sec of mixing with substrate, electron density maps show formation of the covalent thioimidate intermediate at Cys101 in G150T ICH (Fig. 3b, Fig. S7). The substrate selects the native-like, catalytically competent Cys101 conformation exclusively, with no evidence of thioimidate formation on the second Cys101 conformation (Fig. 3b). The absence of thioimidate electron density on the second Cys101 conformation indicates both that the alternate conformation of Cys101 is catalytically inert and that Cys101 stops sampling the alternate conformation once the catalytically competent Cys101 conformation reacts with p-NPIC to form the thioimidate. The refined occupancy of the thioimidate is 0.59, which is less than the 0.85 occupancy of the Cys101 conformer to which it is bonded, indicating partial formation of thioimidate after 30 s. Therefore, the catalytically competent Cys101 conformation is 69% modified to the intermediate in this structure. G150T ICH provides a clear example of catalytic modification of the cysteine nucleophile causing changes to active site conformational dynamics and affords a high-resolution view of combined conformational and chemical heterogeneity in an enzyme active site.

**Figure 3:**
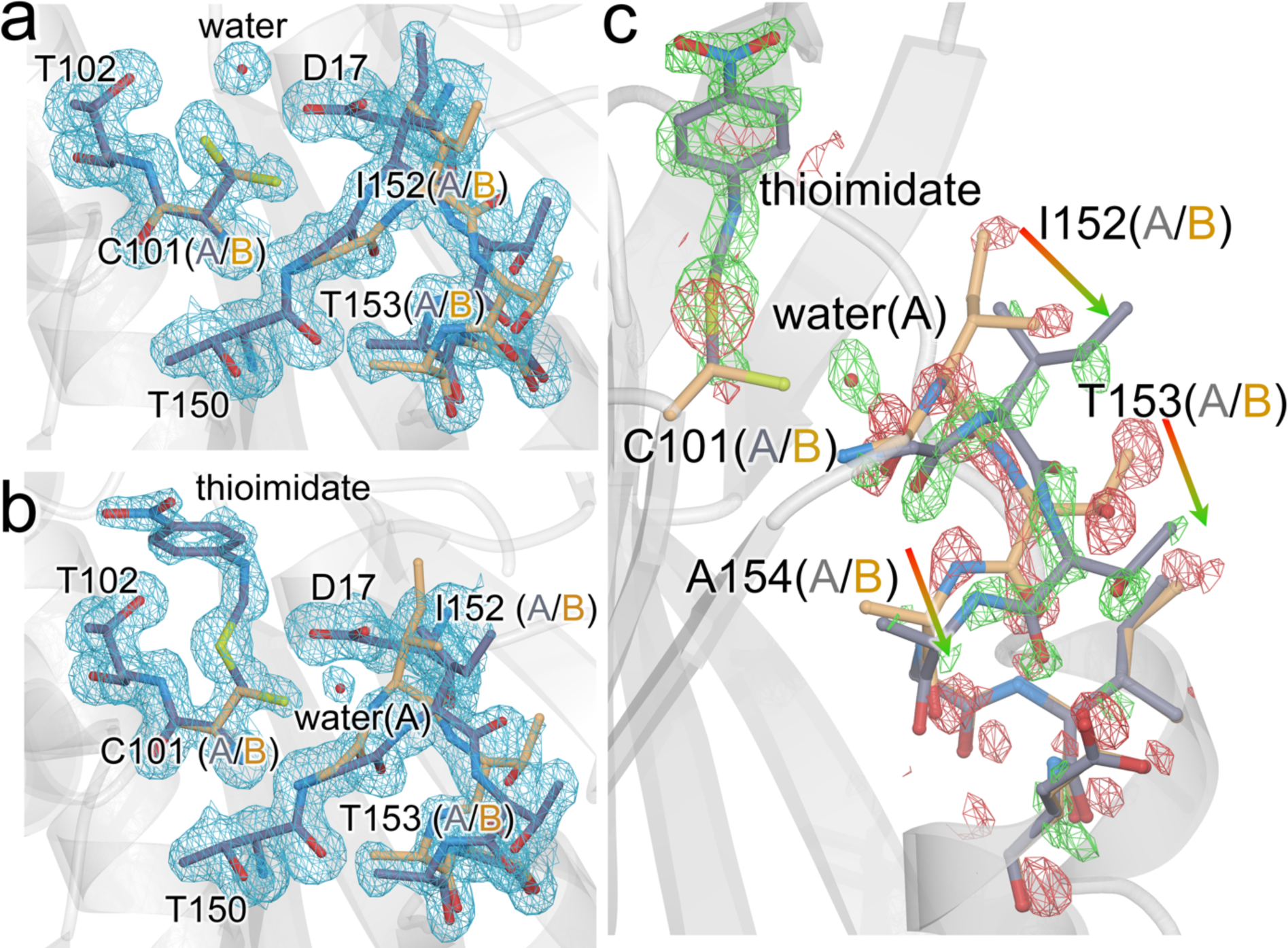
Formation of thioimidate intermediate selects catalytically competent active site conformations and redistributes the G150T ICH ensemble. (a) 2mF_o_-DF_c_ XFEL electron density contoured at 0.9 σ (blue) for the free G150T ICH at 298 K. Alternate conformations for active site residues are blue (conformer A) and gold (conformer B). (b) 2mF_o_-DF_c_ XFEL electron density contoured at 0.9 σ (blue) for the thioimidate intermediate observed 30 seconds after mixing microcrystals with p-NPIC substrate in a mix-and-inject serial crystallography (MISC) experiment. Only conformer A of Cys101 reacts with the intermediate, which redistributes the conformational disorder in the I152 loop. (c) Redistribution of the populations of the active site ensemble upon formation of the thioimidate intermediate. The F_o_(30s)-F_o_(0s) isomorphous difference electron density map is contoured at 3 σ (green, positive/red, negative). Arrows show the direction of population shift, from red (loss of population) to green (gain of population). A partially occupied water (red sphere) enters the active site upon formation of the intermediate.

Formation of the intermediate results in strong F_o_(30s)-F_o_(0s) isomorphous difference electron density extending from the active site to surrounding residues (Fig 3c). The loop comprising residues 151-154 samples two conformations in G150T (Fig.3c), with the one nearer to Cys101 (conformer A in the free enzyme, conformer B in the thioimidate) being more populated when the enzyme is at rest. This will be called the closed conformer. The F_o_-F_o_ difference electron density indicates that the populations of the two loop conformations have been redistributed upon intermediate formation although their structures have not changed significantly (Fig. 3c). The refined occupancy of the closed conformer is reduced from 0.65 in the resting enzyme to 0.39 in the G150T-thiomidiate structure, representing an inversion in the population distribution of these two conformers compared to the free enzyme. However, because the occupancy of the thioimidate is only 0.59, it is likely that a fully occupied catalytic intermediate would result in an even larger reduction in the occupancy of the closed conformer. From a thermodynamic perspective, these two conformers represent local energy minima on the energy landscape of possible G150T ICH configurations, with their relative populations determined by the Boltzmann distribution. The observed change in occupancies between the resting and thioimidate G150T ICH structures shows that thioimidate formation changes the underlying energy landscape such that the relative difference free energy (ΔG) has increased markedly to favor the open conformer. The redistribution of the populations of these two loop conformations also changes the locations of ordered waters around residues 152-154 in helix H and in the surrounding IJ linker region from the other protomer (Fig. S8), demonstrating correlated protein-solvent conformational reorganization. This mobile loop contains Ile152, whose backbone amide donates an H-bond to the Cys101 thiolate and is a severe Ramachandran outlier in the wild-type enzyme. If that H-bond is weakened, Ile152 samples more favorable backbone torsion angles and facilitates the helix shift observed in the wild-type enzyme at low pH (see above) or during catalysis ^32^. In G150T ICH, Ile152 is in Ramachandran-permitted regions in both loop conformers. Therefore, this additional level of loop-helix disorder in G150T ICH was unexpected as it does not appear to be coupled to backbone torsional strain.

### Formation of the catalytic intermediate results in changes to G150T ICH that span active sites in the dimer

Catalysis-activated motions in ICH extend across the dimer interface, spanning both protomers of the ICH dimer. F_o_(30s)-F_o_(0s) isomorphous difference electron density features are distributed throughout the ICH dimer in an anisotropic manner, with most peaks being clustered near the helix H and stretch of residues between helices I and J (IJ linkers) that span the two active sites (Fig. 4a,b). We note that G150T ICH crystallizes in space group C2 with one protomer in the asymmetric unit, meaning the symmetry of these features is a consequence of crystallographic symmetry. However, asymmetry in ICH dynamics typically results in the full dimer being contained in the ASU, as seen in wild-type ICH ^32^. We propose that the fully shifted helix H in G150T ICH causes a symmetrization of ICH protomer dynamics that is obeyed during catalysis in the crystal.

**Figure 4:**
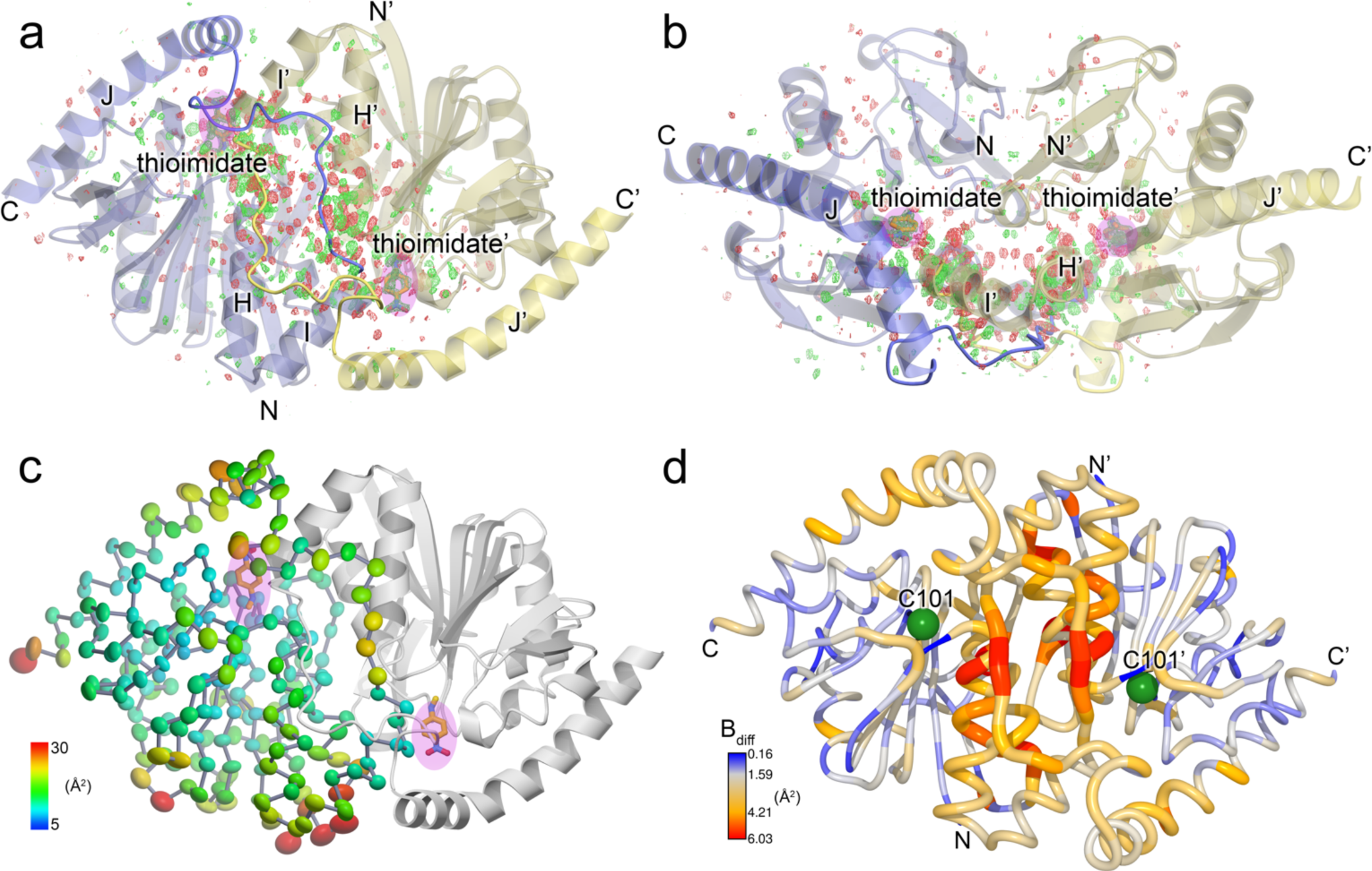
Catalysis causes global changes to G150T ICH dimer conformational ensemble. (a) and (b) Formation of the thioimidate intermediate (labeled; pink highlight) causes changes that propagate across the dimer interface. Two views of the G150T ICH dimer related by a 90° rotation about the horizontal axis are shown. The F_o_(30s)-F_o_(0s) isomorphous difference electron density map is contoured at 3 σ (green, positive/red, negative). The linker connecting helices I and J is shown in darker colors and mediates inter-protomer dynamics that connect the active sites. (c) Anisotropic ADP ellipsoids for Cα atoms at the 85% probability level are colored by magnitude, from blue (5 Å^2^) to red (30 Å^2^). The thioimidate intermediate (labeled; pink highlight) is near areas of elevated mobility. (d) Formation of the thioimidate intermediate increases mobility in several areas of G150T ICH including the IJ linker and helix H. Difference ADP values (B_diff_) between the thioimidate structure and the free enzyme are shown, with scale indicated at the lower left. Areas with higher B_diff_ values coincide with peaks in the F_o_(30s)-F_o_(0s) isomorphous difference electron density map (a, b).

Prior analysis of correlated alternate conformations in wild-type ICH indicated that motions in helix H in one protomer are communicated to the other protomer via the proline-rich IJ linker ^32^. The F_o_(30s)-F_o_(0s) isomorphous difference electron density for G150T shows that even after helix H is shifted, these residues remain a key pathway for dimer-spanning correlated motions. Because each IJ linker region makes contacts with both helices H and H’ in the two protomers in the ICH dimer, displacements in the linker provide a dimer-spanning path between the two active sites (Fig. 4a,b). The IJ linker is a distinctive feature of ICH homologs and distinguishes them from other close structural relatives in the DJ-1 superfamily ^37^, suggesting that other ICH homologs may exhibit similar IJ linker-mediated dynamical pathways through the ICH dimer.

ADPs provide another measure of atomic motions in crystals, and the 1.3 Å resolution of these data allow for the refinement of anisotropic ADPs that contain information about directional preferences in atomic motion. Despite their potentially high information content, comparing ADPs between multiple crystal structures is often complicated by idiosyncratic differences between the crystals used to collect the multiple datasets, leading to mismatches in baseline ADP values and misattribution of crystalline disorder at other length scales to individual atomic motions ^58^. These limitations are partially ameliorated by serial crystallography because a single dataset comprises diffraction images from many crystals, diminishing the influence of individual crystal idiosyncrasies on the refined ADPs. The ADPs in the active site, helix H, and IJ linker regions of the G150T ICH thioimidate intermediate are higher than in the resting enzyme, and these differences correspond closely to the spatial distribution of F_o_-F_o_ difference map features (Fig. 4). These differences are localized primarily to these regions of the dimer, with similar ADPs in other portions of the protein (Fig.4d, Fig. S9). Despite the differences in anisotropic ADP magnitudes between the enzyme at rest and during catalysis, there are no major changes in the directionality of the anisotropic ADPs upon thioimidate formation as determined via visual inspection and Rosenfield rigid body analysis ^59^ (Fig. S10). Therefore, formation of the thioimidate intermediate changes the magnitudes but not overall directional preferences of atomic displacements in the active site, helix H, and IJ linker regions of ICH.

### Water entry into active site depends on dynamics activated by intermediate formation

The ICH mechanism requires hydrolysis of the thioimidate intermediate, although water has not been observed in a catalytically plausible location in prior XFEL electron density maps of the wild-type enzyme during catalysis. By contrast, the G150T ICH thioimidate electron density has a peak consistent with water between Cys101 and Ile152 and 4.3Å away from the thioimidate carbon atom in the scissile C-S bond. This water enters the active site when the thioimidate intermediate forms, as shown by a 5.10 positive peak in the F_o_(30s)-F_o_(0s) isomorphous difference electron density map and the absence of any peak in 2mF_o_-DF_c_ maps of the resting enzyme (Fig. 3). The water is only partially occupied (refined occupancy=0.64) because its entry into the active site requires motion of the 151-154 loop to relieve a steric conflict between the water and Ile152 Cα and Hα atoms (Fig. 5a). Moreover, this water can enter this site only when Cys101 is in its dominant, catalytically competent conformation because of a steric clash with the minor Cys101 conformation (Fig. 5a). Therefore, correlated disorder at multiple locations in the G150T ICH active site gates access of the water molecule to this binding pocket upon formation of the catalytic intermediate (Fig. 5b). This water molecule is not optimally positioned to initiate catalysis, being both too far from the C-S bond and not optimally positioned for an in-line attack and displacement of the Cys101 S atom (Fig. 5a). Therefore, it is possible that it is not the hydrolytic water. However, it is the closest water to the intermediate in the active site and enters the active site upon intermediate formation, suggesting that it is catalytically relevant. The G150T mutant structure provides a clearer view of this state than would be possible in the wild-type enzyme because the shifted helix H conformation required for water entry is transiently sampled with low occupancy once the intermediate forms in wild-type ICH but is the dominant conformation of G150T ICH.

**Figure 5:**
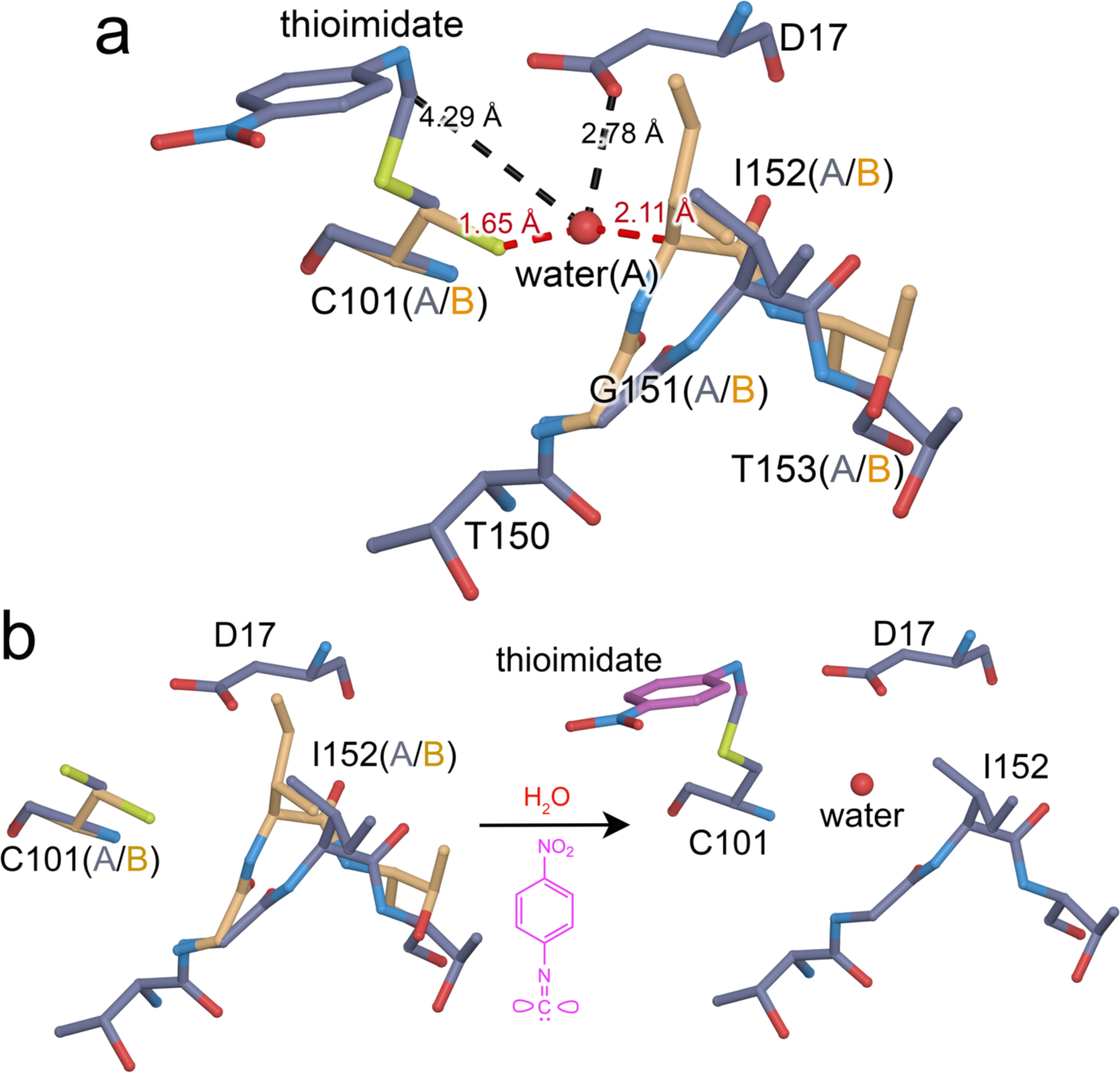
Thioimidate formation and conformational dynamics facilitate water entry into the active site. (a) interactions between the alternate conformations of active site residues and water require correlated motions to avoid steric clashes. The “A” alternate conformation is in slate grey bonds and the “B” alternate conformation is in gold. Interactions between the partially occupied water (labeled red sphere) and surrounding residues are shown with dotted lines, with distances in Ångstroms. Red lines indicate steric clashes between the water and residues in the “B” conformation that are avoided by correlated sampling of the non-conflicting alternate “A” conformations. (b) shows changes in the conformational heterogeneity of the active site upon thioimidate formation and water entry. Reaction of Cys101 with p-NPIC (purple) forms a thioimidate intermediate (purple) and allows water to enter the active site. If both the thioimidate and water are fully occupied, conformational heterogeneity in the active site is reduced to avoid steric clashes.

### Crystalline molecular dynamics simulations show active site dynamics are coupled to Asp17 deprotonation

Asp17 is catalytically essential in ICH, where it is proposed to act as a general acid/base (Fig. 1a) ^37^. In wild-type ICH, Asp17 is protonated (i.e. a carboxylic acid) in the free enzyme as determined using X-ray bond length analysis ^37^, but the 1.3 Å resolution of the XFEL G150T data are too low for this type of analysis. Therefore, the protonation status of Asp17 in the thioimidate intermediate-containing enzyme is unknown. Proton donation by Asp17 to the C1 isocyanide-derived carbon atom to form the thioimidate intermediate is proposed to be a key event in catalysis (Fig. 1a) and should change active site electrostatics, which may be important for the catalysis-activated ICH conformational dynamics that we observe. To investigate the influence of Asp17 ionization state on solvent structure and conformational heterogeneity in G150T ICH, we performed molecular dynamics (MD) simulations of several unit cell volumes containing the G150T ICH thioimidate intermediate with either a protonated Asp17 (residue name ASH, corresponding to the carboxylic acid) or a deprotonated Asp17 (residue name ASP, corresponding to the carboxylate) (Methods). The method of performing molecular dynamics (MD) simulations of volumes of the crystal lattice that we employ here is closely related to those applied to study solvent structure ^60, 61, 62^ and conformational variability ^62^.

To analyze the simulations in the context of the experimental data, we use the MD-MX procedure ^62^. In the MD-MX procedure, structure factors and electron density maps are calculated from the ensemble that was sampled in the simulated crystal volume during the MD trajectory ^62^. This computational time- and lattice-averaging is similar to what occurs during X-ray diffraction from a crystal and thus facilitates the direct comparison of the simulation to experiment. We used these simulation trajectory-derived structure factors (called F_MD_ hereafter) to calculate isomorphous difference electron density maps by subtracting the MD structure factors for G150T ICH-thioimidate with a protonated Asp17 (ASH) from the same form of the protein with a deprotonated Asp17 (i.e. F_MD_(ASP)-F_MD_(ASH)) (Fig. 6a). The MD-MX difference map indicates Asp17 deprotonation reorganizes solvent in the active site and redistributes the populations of the alternate conformers in the I152 loop (Fig 6c). Because the Cys101-thioimidate is present in both simulations, Asp17 deprotonation alone is responsible for this change in active site dynamics. Comparison of the F_MD_(ASP)-F_MD_(ASH) difference electron density map (Fig. 6c) to the experimental F_o_(30s)-F_o_(0s) difference map (Fig. 6b,d) shows a remarkable agreement in features, demonstrating that many of the correlated changes in enzyme dynamics and solvent entry in the active site observed in the G150T ICH XFEL experiment can be explained by the deprotonation of Asp17. Therefore, MD-MX analysis supports the mechanism in Fig 1a and demonstrates that conformational heterogeneity in the ICH active site is highly sensitive to the protonation state of Asp17, which changes during catalysis. In combination with the previously demonstrated importance of Cys101 ionization for the initial helix H displacement ^32^, these results underscore the tight coupling between active site electrostatics and enzyme dynamics in ICH.

**Figure 6:**
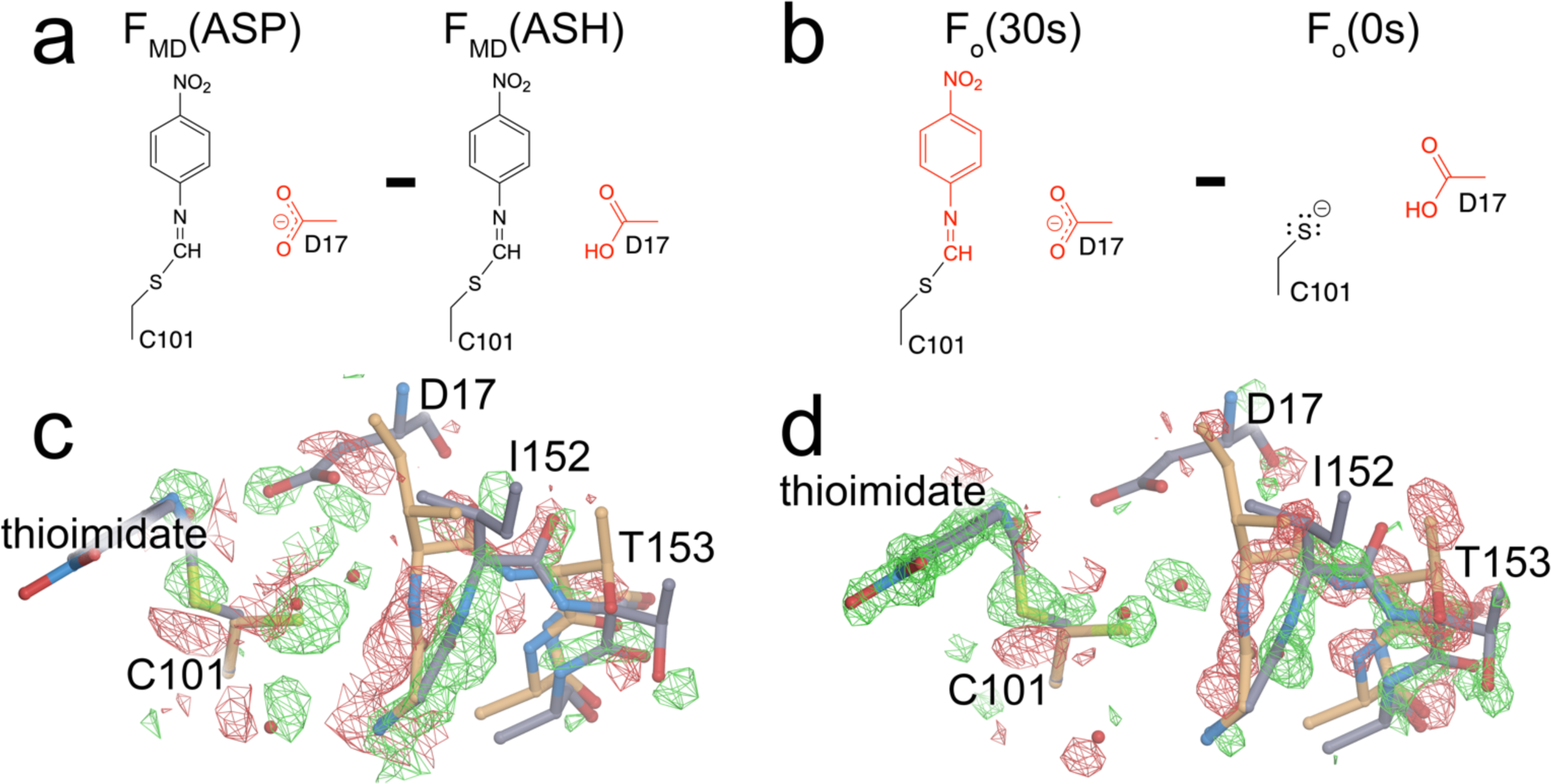
Deprotonation of Asp17 causes major conformational changes in the active site. (a) shows the differing protonation states of Asp17 that were used to calculate structure factors (F_MD_) in crystalline MD simulations. (b) shows the corresponding state of the enzyme active site in the experimental F_o_(30s)-F_o_(0s) isomorphous difference electron density map. (c) shows the calculated isomorphous difference electron density map described in (a) contoured at 6 α and (d) shows the experimental difference electron density map for the active site contoured at 3 α. The excellent agreement between these maps (c,d) demonstrates that water entry and changes in the active site conformational ensemble upon thioimidate formation are driven by deprotonation of Asp17.

## Discussion

Mix-and-inject serial crystallography (MISC) allow crystalline enzymes to be observed during catalysis, opening a window into non-equilibrium conformational changes that have been difficult to observe using other approaches. The 1.3 Å resolution of these data are unusually high for room-temperature XFEL serial crystallography and permit a detailed modelling of conformational disorder as well as the refinement of anisotropic ADPs. We find that introduction of the substrate selects catalytically competent conformations of the G150T ICH active site and that there are pathways of correlated atomic motion that extend across the enzyme dimer. Interpreting our results from an energy landscape perspective ^4^, binding of substrate to the G150T active site changes the depths of pre-existing energy minima, thereby altering populations of the conformers in the ensemble in the absence of large overall structural changes and favoring a subset of possible trajectories along the energy surface (Fig 7a,b). We note that it is difficult to model conformations with populations below approximately 5-10% occupancy in X-ray crystallography, so it is possible that some conformations are sampled but insufficiently occupied to be modeled. Bearing this limitation in mind, the resting G150T ensemble contains the major conformations observed upon formation of the thioimidate intermediate, while wild-type ICH populates distinct conformations during catalysis in response to covalent modification of Cys101 ^32^. Because the G150T mutation favors conformations that are sampled at low occupancy by the wild-type enzyme during catalysis, it provides an example of the value of using mutagenesis to increase the populations of regions of conformational space that are difficult to observe in wild-type enzymes.

**Figure 7:**
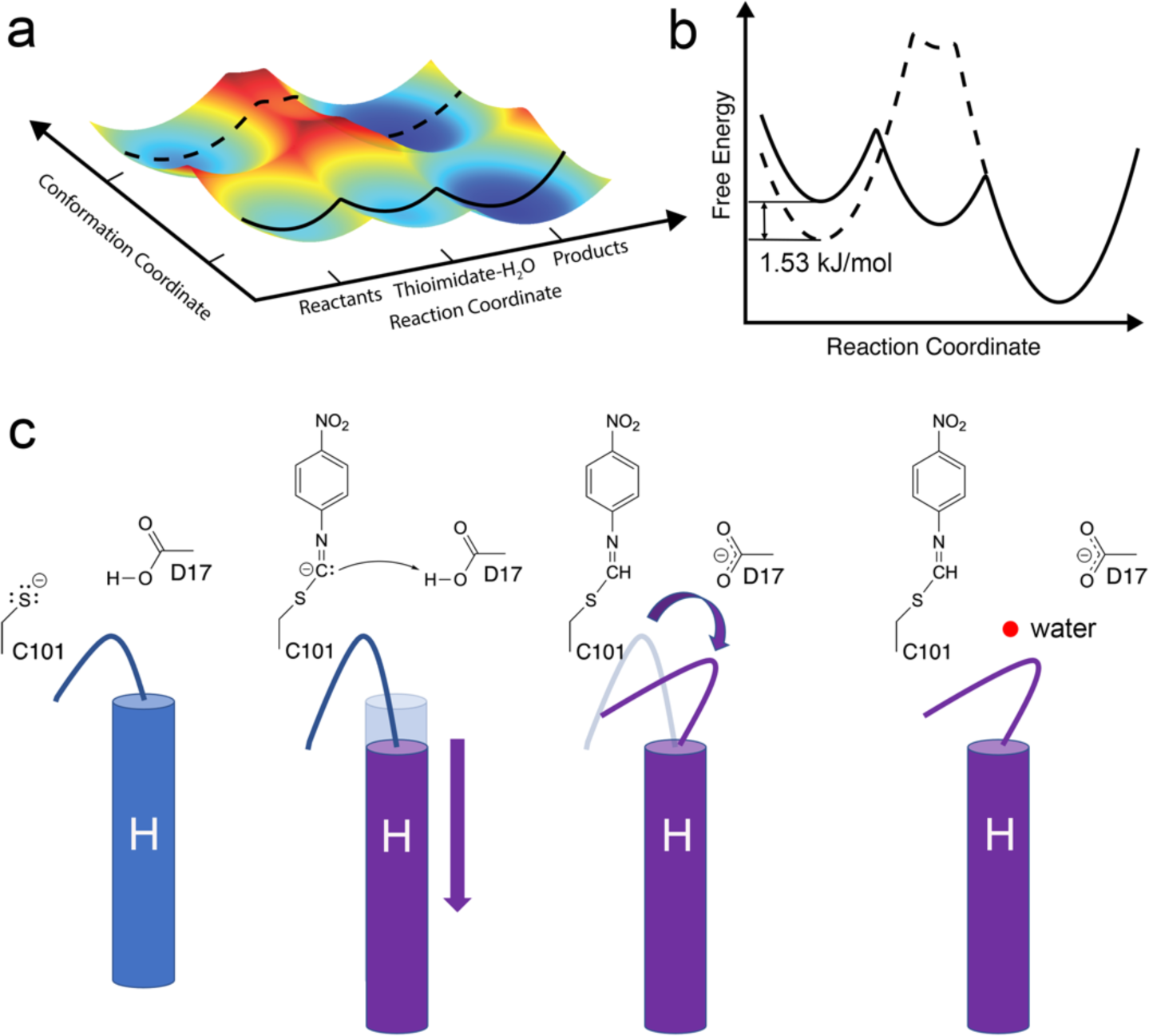
Energy landscape model of ICH catalysis and charge-coupled enzyme dynamics. (a) shows a plot of free energy of the two active site loop conformations vs. progress along the reaction coordinate as inferred from XFEL structures. As the reaction proceeds through the intermediate, the open loop conformation (bottom, with solid line) is energetically preferred. (b) slices through the surface are show in a convention two-dimensional free energy vs. reaction coordinate plot. The 1.53 kJ/mol energy difference of the loop conformations in the resting enzyme calculated from the crystallographic occupancies using the Gibbs free energy equation (see Methods). The closed loop conformation (dotted line) has a higher energy than the open one upon intermediate formation and concomitant Asp17 ionization. (c) A schematic of catalysis in ICH. Neutralization of charge on Cys101 triggers an initial shift in helix H, followed by ionization of Asp17 causing a change in loop position that allows water (red sphere) to enter the active site.

Previous pre-steady state kinetic results indicate that the G150T mutation slows steps both before and after thioimidate formation ^32^. The slowness of G150T was an asset for this study, as it allowed for long substrate mixing times (30s in our case) and accumulation of the thioimidate intermediate and active site water molecule. It is tempting to ascribe the catalytic impairment of G150T to the conformational disorder we observe at Cys101. However, the ∼0.2 refined occupancy of the catalytically incompetent Cys101 rotamer does not explain the ∼80% decrease in k_cat_ (steady-state) or the ∼60% decrease in the pre-steady state burst rate constant for G150T ICH compared to the wild-type enzyme ^32^, indicating that there are other contributors to this mutant’s catalytic impairment. In our view, the lost H-bond interaction between the Cys101 thiolate and the backbone amide of Ile152 is likely to be an important contributor to G150T ICH’s slower kinetics, as this H-bond should affect the Cys101 pK_a_ value, may indirectly influence the extent of Asp17 protonation, and potentially stabilize the thiolate leaving group during hydrolysis of the thioimidate intermediate.

Despite having nearly identical overall structures, isomorphous difference (F_o_-F_o_) electron density maps calculated between the 274K synchrotron and 298 K XFEL datasets for G150T ICH showed features that aligned with principal axes of anisotropic ADPs and with minor shifts in atomic position between these higher temperature structures and a cryogenic (100 K) structure. The overlap between these F_o_-F_o_ electron density peaks and modes of intrinsic mobility in the protein leads us to propose that the difference in temperature is most likely responsible for these changes. Recognizing that many aspects differ between single crystal synchrotron and serial XFEL data collection, we cannot definitively determine the cause of these F_o_-F_o_ electron density features. However, our attribution of these map features to temperature-induced changes is supported by prior work using multiple temperature and temperature-jump crystallography, which concluded that proteins respond to modest changes in temperature with complex, distributed changes in their conformational ensembles ^27, 54^. The sensitivity of isomorphous difference maps to moderate changes in temperature (24°C in this case) should be borne in mind when interpreting time-resolved crystallography studies where significant temperature changes are possible, such as heating during photoinitiation using high-intensity lasers, evaporative cooling during liquid jet sample delivery in vacuum, or mixing liquids of differing temperatures.

The entry of a water into the G150T ICH active site upon thioimidate formation is intriguing, as it offers a view of an important step prior to intermediate hydrolysis. Although we cannot be certain that this water is the hydrolytic water, it is the first time that water has been observed entering the ICH active site during catalysis and is likely to be functionally important. Movement of the I152 loop in the shifted helix H conformation is required for water to bind at this location, explaining why this water molecule was not observed in the wild-type enzyme-thioimidate structure, where the helix is predominantly in the unshifted conformation ^32^. In contrast, helix H is constitutively shifted in G150T ICH-thioimidate and therefore this rare event in the wild-type enzyme is sampled enough to permit modelling in G150T ICH. Comparison of experimental and MD-MX electron density maps shows that the deprotonation of Asp17 is a major driver of the additional loop motion and water binding in the active site upon thioimidate formation. The MD-MX result explains why the crystal structures of wild-type ICH at low pH do not show electron density supporting either the additional 152 loop motion or the water. Although these wild-type ICH structures have a shifted helix H due to Cys101 protonation, Asp17 is also protonated. By contrast, the postulated charge state of the active site containing the thioimidate intermediate is a neutral sulfur atom and a deprotonated (carboxylate) Asp17 (Fig. 1a), which cannot be experimentally recapitulated by manipulating pH alone. These results highlight the value of using MD-MX to study changes in enzyme conformational dynamics that are coupled to active site electrostatics, because directly determining ionization states is extremely difficult using macromolecular X-ray crystallography alone. MD-MX provides a powerful means to manipulate ionization states in silico and then compare the structure factors calculated from these simulations directly to time-resolved experimental data using isomorphous difference electron density maps, as we have done here. The combination of MD simulation and X-ray crystallography experiments suggests a mechanism for how changes in ICH active site electrostatics modulate the enzyme conformational ensemble as a function of catalysis (Fig. 7c).

A challenge in time-resolved crystallography is interpreting electron density maps that result from stochastic conformational changes that are not synchronized among unit cells, leading to a blurring of the electron density and difficulties in its interpretation. The present study demonstrates the value of using mutants to enrich enzyme ensembles in rare conformations that are difficult to study in wild-type enzymes. In combination with experimental interventions that alter enzyme kinetics to favor to accumulation of reaction intermediates, mutational probing of conformational ensembles can extend the utility of time-resolved crystallography to more difficult systems or less kinetically accessible regions of their reaction coordinates. Because ICH diffracts to high resolution, has correlated dynamics that change during catalysis, and exemplifies several mechanistic features common to cysteine-dependent enzymes, it is a valuable model system for developing new approaches for time-resolved structural enzymology.

## Methods

### G150T ICH protein purification, crystal growth, and conventional X-ray data collection

The G150T mutant of *Pseudomonas fluorescens* isocyanide hydratase (PfICH) G150T was purified as previously described ^32^. Briefly, 1.5L cultures of LB were inoculated and grown to 0.4 OD_600_ at 37°C. Protein expression was induced with 200 uM IPTG for four hours at 37°C, followed by cell harvest by centrifugation, resuspension in lysis buffer (50 mM HEPES pH=7.5 and 300 mM NaCl), lysis by sonication, and centrifugation to remove cell debris. The supernatant was passed over Ni^2+^-NTA resin (Sigma) and the protein was eluted with 250 mM imidazole pH=7.0 in lysis buffer. The N-terminal 6×His tag was removed by cleavage with thrombin during dialysis into against 25mM HEPES pH=7.5, 150mM KCl at 4°C. Purified protein was then concentrated to 40mg/mL by centrifugal concentration (Millipore).

G150T ICH crystals used to obtain the cryogenic structure were grown using sitting drop vapor diffusion with 23mg/mL protein against a reservoir of 23% PEG 3350, 200 mM MgCl_2_, 100 mM Tris-HCl pH=8.8, and 2 mM DTT as previously reported ^32^. These crystals were cryoprotected by serial transfer through the reservoir solution supplemented with increasing concentrations of ethylene glycol to a final concentration of 30% v/v and then plunge-cooled in liquid nitrogen. The crystal structures of wild-type ICH at various pH values were grown by hanging drop vapor equilibration at 20°C by mixing 2 μl of 20mg/mL wild-type ICH protein with 2 μl of reservoir solution. Reservoir solutions contained 23% PEG 3350, 200 mM MgCl_2_, 2 mM DTT and 100mM of either Tris-HCl, (pH=8.3) sodium citrate (pH=6.0), or sodium acetate (pH=4.2, 5.0, or 5.4). Crystals grew in 2-7 days and were cryoprotected and plunge-cooled as described above. Crystal growth, mounting, and data collection for the 274 K G150T ICH synchrotron data was previously described ^46^. In that prior study these data were indexed in an alternative setting in space group I2 for compatibility with a diffuse scattering workflow but were reindexed in space group C2 for this work, permitting calculation of isomorphous difference electron density maps with the XFEL data.

Single-crystal diffraction data were collected at beamlines 12-2 and 9-2 at the Stanford Synchrotron Radiation Lightsource (SSRL) using the oscillation method with shutterless data collection and a Dectris Pilatus 6M pixel-array detector. The wild-type ICH pH series and the cryogenic crystal structure of G150T ICH were collected at 100 K in nylon loops, while the triplicate room-temperature G150T synchrotron datasets were collected in 10 μm thick glass number 50 borosilicate capillaries glass capillaries (Hampton Research) at 274 K as previously described ^46^. The data were indexed and integrated using XDS ^63^ and scaled using Aimless ^64^. Data statistics are presented in Table S1.

### Microcrystal growth, mix-and-inject serial crystallography, XFEL data collection and processing

Microcrystals of G150T ICH were grown by seeding. 100-150 G150T ICH crystals measuring ∼100 μm x 50 μm x 50 μm were harvested in stabilizing solution (15.5% PEG 3350, 125 mM MgCl_2_, and 100 mM Tris-HCl pH 8.8) and pulverized by vortexing for ≥10 minutes with approximately 50 0.5mm stainless steel beads (Next Advance). The pulverized crystal seed stock was decanted and then centrifuged at 84×g for 1 min, decanted again, and centrifuged a second time at 325×g for 1 min to remove larger uncrushed crystal fragments and other detritus. A 1:200 dilution of this seed stock in crystal growth solution (31% PEG 3350, 250 mM MgCl_2_, and 125 mM Tris-HCl pH 8.8) was then mixed with an equal volume of purified G150T PfICH (40mg/mL in 25 mM HEPES pH 7.5, 100 mM KCl) and gently mixed by inversion. G150T ICH microcrystals of approximate dimensions 30 μm x 10 μm x 10 μm grew within 30 minutes and further growth was quenched by adding 0.5 mL of 1.15× stabilizing solution per 1 mL of crystals. Crystals were stored at room temperature and used within five days after growth.

XFEL serial crystallography data were collected at the Macromolecular Femtosecond Crystallography (MFX) endstation of the LCLS ^65^ during beamtime mfxlx4418. XFEL pulses at 12 keV with ∼1×10^12^ photons per 40 fs pulse were delivered to the interaction region at 30 Hz repetition. The X-ray spot size at sample was ∼3 μm in diameter. Diffraction data was collected on a Rayonix MX340-XFEL CCD detector operated in 4×4 binning mode and hits were analyzed in real time using OnDA ^66^. The powder diffraction pattern from silver(I) behenate was used to estimate the detector position and the location of the beam center. Joint refinement of the crystal models were performed against the detector position for each batch to account for small time-dependent variations in detector position^2^.

The concentric-flow microfluidic electrokinetic sample holder (coMESH) injector ^67^ was used to deliver G150T ICH microcrystals to the XFEL beam at room temperature and under normal atmosphere composition and pressure. The sample was held in a custom stainless steel sample reservoir that was agitated during the experiment to prevent crystal settling. A Shimadzu LD20 HPLC pump hydraulically actuated a teflon plunger to advance the sample slurry. A 100 μm x 160 μm x 1.5 m fused silica capillary (Molex) connected the reservoir and filters to the coMESH. This capillary continued unobstructed through the center of the microfluidic tee (IDEX-HS) and into a concentric 250 μm x 360 μm x 500 mm capillary. The capillaries were optically aligned to obtain an approximate 0 mm offset for the free enzyme (0s) dataset, then were recessed and measured externally to achieve the 62 mm recess for the 30s substrate mixing dataset. The outer annulus of the coMESH flowed the stabilizing solution through the perpendicular branch of the microfluidic tee in the case of the free dataset and flowed saturated (∼2-3 mM) p-NPIC substrate in stabilizing solution for the 30s timepoints. The p-NPIC substrate was synthesized as previously described ^32^. DTT was omitted from all solutions because it reacts with the p-NPIC substrate. The p-NPIC substrate was loaded into a similar stainless steel reservoir as the crystal slurry and attached to 250 μm x 1/16” x 1 m PEEK tubing (Zeuss) connected to the side of the coMESH microfluidic tee junction. Like the crystal slurry, the fluid was actuated by a second HPLC pump (Shimadzu) at 3 μl/min. The 1 m of PEEK tubing was interrupted by a stainless steel union (IDEX-HS) that was electrically charged at +3.1 kV (Stanford Research Systems, SRS PS300). This imparted an electrical charge to the wetted fluid and ultimately focused the ensuing meniscus of mixed sample and stabilizing solution towards the interaction point of MFX. The outer liquid focused the fluids ∼0.5 mm away from the outer capillary towards the XFEL focus at the Macromolecular Femtosecond Crystallography (MFX) endstation. The charged meniscus was approximately 5 mm away from a grounded counter electrode to complete the electrokinetic focusing. The time delays were assumed to be sufficiently long as compared to the electrokinetic mixing phenomena and were determined by the time the bulk fluid would traverse the offset distance. The flow rates and voltages were held constant during each time point. The flow of the crystals through the main sample capillary (0.1 x 0.16 x 1500 mm) were optically monitored with a 50x objective to assure minimal flow deviations, in addition to the backing pressure of the pumps driving the flows. The stabilizing solution was not introduced to the crystal-carrying slurry until the samples were 62 mm away from the X-ray interaction point. This constraint, coupled with combined volumetric flow rates of 6 μl/min (3 µl/min for the crystal slurry + 3 µl/min for the stabilizing solution with or without substrate), dictated the delay times. Approximately 500 µl of microcrystalline sample was used to measure each timepoint.

Serial crystallography datasets were reduced and processed using *cctbx.xfel* and DIALS ^68, 69^. Data were scaled and merged to their final resolution cutoffs using *cctbx.xfel.merge* with errors determined by the ev11 method ^70^. For the free enzyme data, 23990 had their intensities integrated and merged. For the 30 s thioimidate data, 22042 frames had their intensities integrated and merged. The high resolution cut-off values for the final datasets were determined by previously established resolution cutoff criteria, such as ∼10× multiplicity, the point where the I/σ(I) values become stable, and where CC_1/2_ values stop decreasing monotonically ^71^, indicating no useful information is contained in resolution shells beyond that point. The data statistics for the XFEL data are provided in Table S2.

### Model refinement and analysis

All models were refined in PHENIX ^72^ using riding hydrogen atoms against a maximum likelihood target based on structure factor intensities. Individual anisotropic ADPs were refined for all models except wild-type ICH at pH=4.2 and 5.0, where the translation-libration-screw (TLS) model was used with automatically determined rigid body boundaries in PHENIX. Stereochemical and ADP weights were optimized. Initial G150T ICH XFEL models were based on PDB 6NI4 ^32^ and were manually adjusted in Coot ^73^ and validated using tools in Coot and MolProbity ^74^. Chloride ions were assigned based on the magnitude of the 2mF_o_-DF_c_ electron density peaks and the presence of weak anomalous difference electron density in the cryogenic G150T ICH structure. Alternate conformations were manually built and refined with grouped occupancies when a contiguous stretch of residues was modeled in alternate positions. However, the thioimidate intermediate only partially modifies Cys101 in conformer A, therefore these occupancies are unequal and a result of a combination of conformational and chemical heterogeneity. The previously collected 274 K synchrotron G150T ICH data were reindexed in C2 and refined as described above. All refined model statistics are in provided in Table S3.

The difference in ADPs (B-factors) between the 30 second and 0 second G150T ICH XFEL models were calculated for the main chain atoms and then averaged for each residue using Baverage in the CCP4 suite ^75^. For residues in alternate conformers, the altloc A conformer (which is typically the most occupied conformer) was used for this calculation. Rosenfield difference matrices ^59^ of main chain atoms were calculated in ANISOANL in the CCP4 suite ^75^ using 30 bins. The free energy difference between the two 151-154 loop conformations in the free enzyme were calculated using the ratio of their refined occupancies and the Gibbs free energy equation: 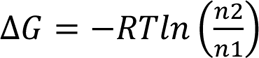, where R is the ideal gas constant, T is temperature in K (298 K in this experiment), and n_i_ is the occupancy of the i^th^ conformer. All isomorphous difference maps were calculated in PHENIX with no weighting. The pH dependence of helix H conformations in Fig. 3b was fitted using the Henderson-Hasselbalch equation: 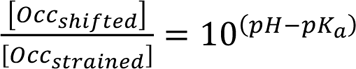

### Rate vs. pH profile of wild-type ICH

The pH-dependence of wild-type ICH catalysis was measured using an acetic acid/sodium phosphate double buffer with pH values in the range of 3.5-10.25. Solutions were prepared by adjusting the pH of a 2 mM acetic acid and 2 mM sodium phosphate (monobasic) double buffer with small volumes (0.5-1 µL) of 500 mM NaOH. Catalysis was initiated by the addition of 0.5 µM wild-type ICH to the double buffer containing 100 µM p-NPIC substrate and monitored by absorbance at 320 nm for 60 seconds using a Cary50 spectrophotometer. Formation of the product (para-nitrophenyl formamide) was linear in time for all measurements and the slope of the best-fit line (A_320_/sec) at each pH was converted to units of µM product/sec using an extinction coefficient of 1.33×10^4^ M^-1^ cm^-1^ for the product ^32^. After the assay was complete, the pH of each sample was measured at its working concentration using an Orion PerpHecT ROS pH microelectrode (Thermo Scientific).

### Crystalline Molecular Dynamics Simulation

Three different crystalline molecular-dynamics (MD) systems were prepared, based on the free enzyme and Cys101-thioimidate intermediate G150T ICH crystal structures. In one system, the free enzyme crystal structure was used to construct a 2×2×2 unit cell model of the protein in the crystalline state. Monte Carlo sampling was used to determine the conformation of multi-conformer residues for each protein according to the occupancies in the crystal structure. In the other two systems, the thioimidate intermediate crystal structure was used to construct a 2×2×2 unit cell model of the protein in the crystalline state: in one system, Asp17 was modeled as unprotonated (ASP17) whereas in the other system, Asp17 was modeled as protonated (ASH17). In both systems, Monte Carlo sampling was used to determine the conformation of multi-conformer residues for each protein. However, for the thioimidate intermediate residue, an additional round of Monte Carlo sampling was used to determine whether Cys101 in the conformation consistent with covalent modification would have the adduct appended. This was done because Cys101 samples two conformations, only one of which is catalytically active and that conformation (altloc A) is only partially modified to the thioimidate. In all cases, *cctbx* ^72^ methods were used to read in the crystal structure .pdb files and propagate each protein in the state determined by Monte Carlo sampling to a different position in the supercell system. The systems were prepared, solvated, and neutralized with ions using GROMACS version 2022.4 ^76^ methods (*pdb2gmx*, *solvate*, and *genion,* respectively). In each system, additional Cl^-^, K^+^, and Mg^2+^ ions were added, to best reproduce the concentrations of the crystal buffer (125mM MgCl_2_, 67 mM Tris-HCl, 12 mM HEPES, 50 mM KCl). The protein volume was calculated with the ProteinVolume Online server from *gmlab.bio.rpi.edu* ^77^, and the total volume of the 32 proteins (8 unit cells, with 4 proteins in each) was subtracted from the volume of the supercell, to arrive at a solvent volume of 1.0379×10^6^ Å^3^ or 1.0379×10^-21^ L. This solvent volume required an additional 230 Cl^-^ ions, 78 Mg^2+^ ions, and 74 K^+^ ions, on top of the ions required for neutralization (80 K^+^ ions for the apo and ASP17 system, 112 K^+^ ions for the ASH17 systems). The protein and ions were parameterized with the AMBER14SB ^78^ forcefield and the waters were parameterized with the TIP3P parameter set. The parameters for the Cys101 thioimidate intermediate (CYT) residue were determined using *MRP.py* ^79^, a software package for parametrization of post-translationally modified residues, which uses AmberTools ^80^ and Gaussian 16 ^81^ to determine the bonded and non-bonded parameters and partial charges.

Supercell systems were subjected to iterative rounds of solvation and equilibration using *gmx solvate* and *gmx mdrun* to bring the systems up to pressure in the constant particle number, volume, and temperature (NVT) ensemble (at 300K). In contrast to the more common NPT ensemble, in which the side of the simulated box are adjusted to tune the pressure, NVT ensembles are necessary for supercell systems to maintain the correct crystalline symmetry and crystal contacts. 100ns of production simulation was performed for all systems, with time steps of 2 fs; neighbor searching was performed every 10 steps; the PME algorithm was used for electrostatic interactions with a cutoff of 1 nm; a reciprocal grid of 96 x 80 x 72 cells was used with 4^th^ order B-spline interpolation; a single cut-off of 1 nm was used for Van der Waals interactions; temperature coupling was done with the V-rescale algorithm. Harmonic positional restraints, with a restraint constant of 200 kJ^-1^mol^-1^nm^-2^ were applied to all heavy atoms in the system, with the initial propagated crystal-structure supercell positions as the target. The root-mean-squared deviation (RMSD) from the initial heavy atom positions was used to monitor the relaxation of the system under the restraints to a steady-state ensemble, with all systems arriving at steady state at around 60ns.

The final 10ns of simulation were used for analysis using the MD-MX procedure introduced in Wych et al ^62^. The *xtraj.py* Python script ^82^ in the LUNUS GitHub repository (https://github.com/lanl/lunus) was used to calculate the simulated structure factors consistent with the 10ns of simulation used for analysis. Using this method, it is possible to compute the structure factors from any component of the system; structure factors were computed for the full system, the protein atoms, the water atoms, and for the atoms of each ion (Mg^2+^, Cl^-^, and K^+^), individually. “Protein-first refinement” ^62^ was performed with PHENIX ^72^ using simulated intensities computed from the structure factors calculated from just the protein atoms, the structure factors computed from the full system, and the intensities measured in experiment, to produce structural models consistent with both the MD and experimental data. For each system, the proteins in the final frame of the simulation were mapped back on to the asymmetric unit with *cctbx* tools using the unit cell and space group information, to serve as a representation of the ensemble present across all the proteins in the crystalline system. In the present study, the procedure was primarily used to produce isomorphous difference maps using the simulated structure factors. The difference maps were computed with *sftools* from CCP4 ^75^. The full MD-MX procedure also was used to produce revised crystal structures ^62^ with the following R factors: R_work_=0.1492, R_free_=0.1885 for the free enzyme; R_work_=0.1584, R_free_=0.1897 for ASP17; R_work_=0.1600, R_free_=0.1989 for ASH17.

## Abbreviations

ADP: atomic displacement parameter
ASU: asymmetric unit
coMESH: concentric-flow microfluidic electrokinetic sample holder
DTT: dithiothreitol
HEPES: 4-(2-hydroxyethyl)-1-piperazineethanesulfonic acid
ICH: isocyanide hydratase
IPTG: isopropyl-β-D-1-thiogalactopyranoside
LCLS: Linac Coherent Light Source
MISC: mix-and-inject serial crystallography
PDB: Protein Data Bank
PEEK: Polyether ether ketone
PEG: polyethylene glycol
p-NPIC: para-nitrophenyl isocyanide
RMSD: root-mean-square deviation
TLS: translation-libration-screw
UV: ultraviolet
XFEL: X-ray free electron laser

## Acknowledgements

Use of the Stanford Synchrotron Radiation Lightsource, SLAC National Accelerator Laboratory, is supported by the U.S. Department of Energy, Office of Science, Office of Basic Energy Sciences under Contract No. DE-AC02-76SF00515. The SSRL Structural Molecular Biology Program is supported by the DOE Office of Biological and Environmental Research, and by the National Institutes of Health, National Institute of General Medical Sciences (P30GM133894). Use of the Linac Coherent Light Source (LCLS), SLAC National Accelerator Laboratory, is supported by the U.S. Department of Energy, Office of Science, Office of Basic Energy Sciences under Contract No. DE-AC02-76SF00515. The contents of this publication are solely the responsibility of the authors and do not necessarily represent the official views of NIGMS or NIH.

## Data Availability

All refined structural models and structure factor data are deposited with the Protein Data Bank (PDB) with accession codes: 8TSF (G150T ICH XFEL free enzyme), 8TSN (G150T ICH XFEL thioimidate intermediate, 30 seconds mixing), 8TSX (G150T ICH 100 K), 8TSU (G150T ICH 274 K synchrotron dataset 1), 8TSY (G150T ICH 274 K synchrotron dataset 2), 8TSZ (G150T ICH 274 K synchrotron dataset 3), 8TT0 (wild-type ICH pH=4.2), 8TT1 (wild-type ICH pH=5.0), 8TT2 (wild-type ICH pH=5.4), 8TT4 (wild-type ICH pH=6.0), 8TT5 (wild-type ICH pH=8.3).

## Author Contributions

MEW, RGH, MCT, AHF, MAW planned the experiments; VKT, NK, DBB generated key reagents; NS, MD, DCW, CD, RGS, SL, DMR, AC, SB, MSH, CK, FP, FRM, ASB, NKS, IDY, AMW, MCT, AHF, MEW, MAW performed the experiments; NS, MD, DCW, ASB, NKS, IDY, AMW, AHF, MEW, MAW analyzed and interpreted the data; DCW, MEW, MAW wrote the manuscript; NS, MD, DCW, CD, RGS, DMR, NKS, IDY, AMW, MCT, AHF, MEW, MAW edited the manuscript.

