## Supporting information for "Changes in an Enzyme Ensemble During Catalysis Observed by High Resolution XFEL Crystallography"

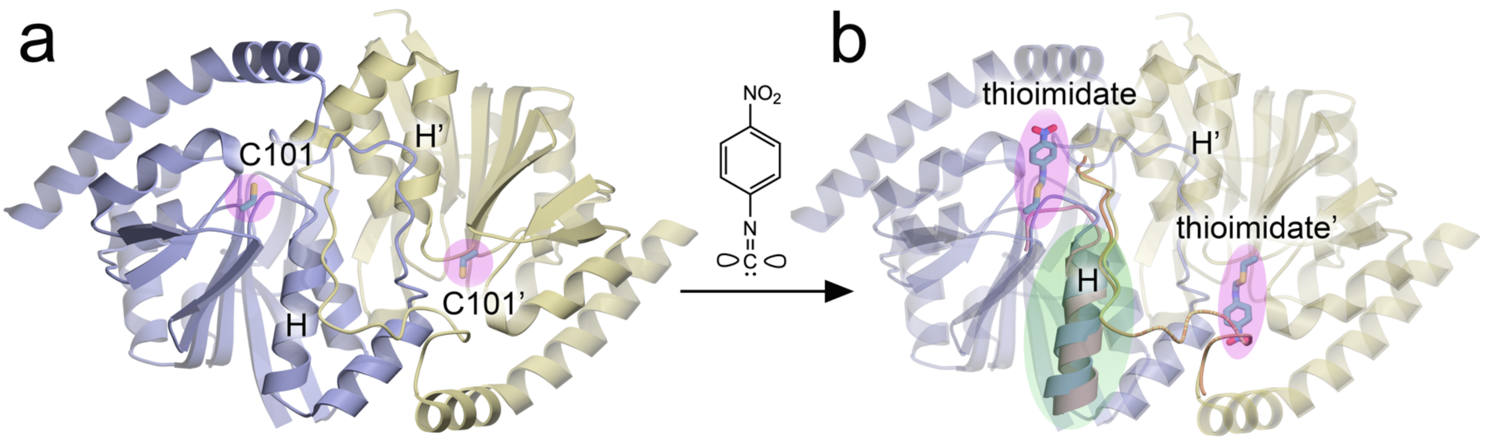


**Figure S1. Schematic of wild-type ICH catalysis-activated motions**. (a) Wild-type ICH is shown in ribbon cartoon with the active site highlighted in pink. (b) Upon formation of the thioimidate intermediate, changes in active site hydrogen bonds allow sampling of a shifted conformation of helix H (green highlight).

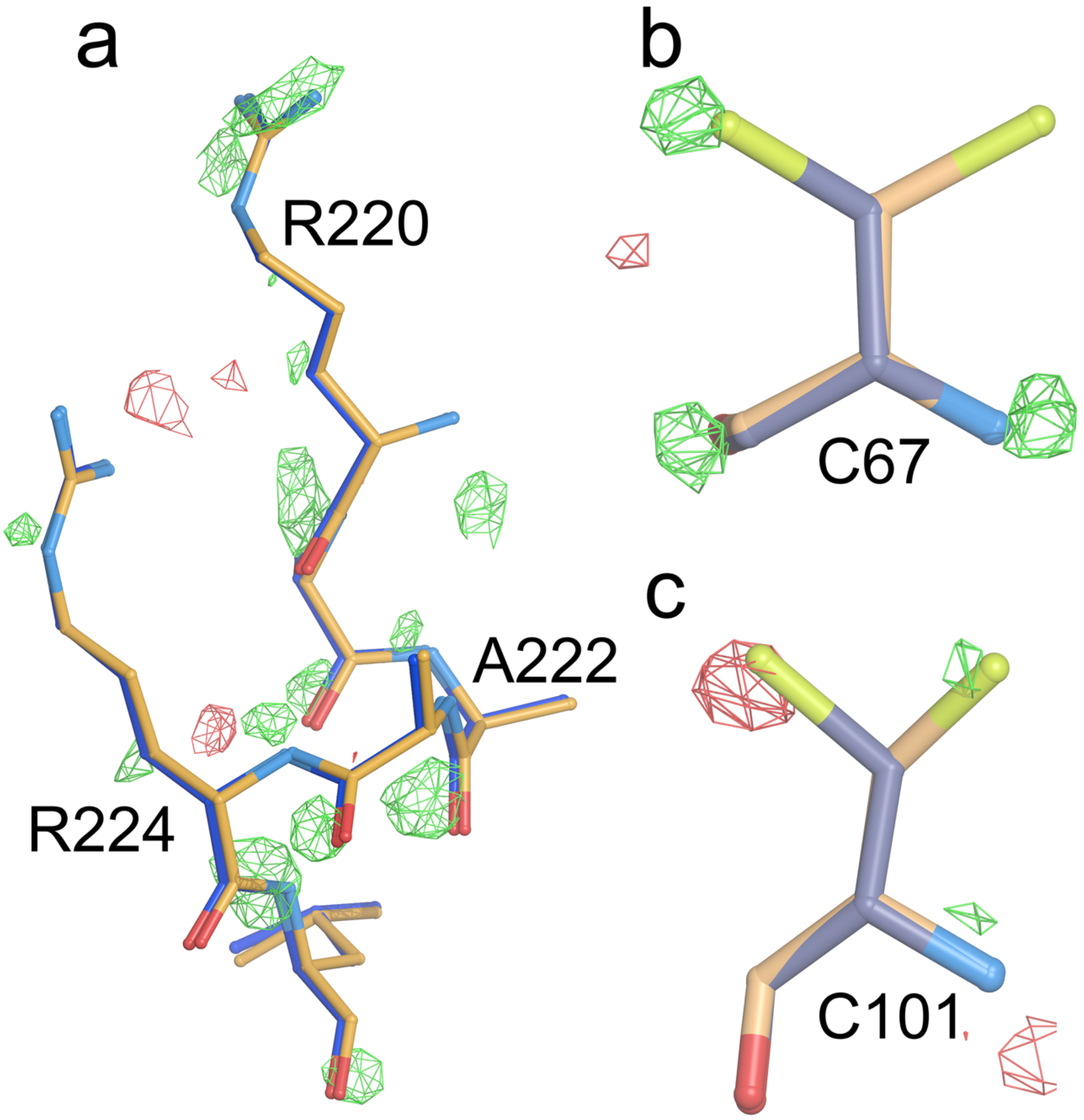


**Figure S2. Examples of temperature-dependent differences in G150T ICH.** $F_o$(274 K synchrotron)-$F_o$(298 K XFEL) isomorphous difference electron density maps contoured at 3 $\sigma$, with positive features in green and negative ones in red. (a) Difference electron density peaks in these residues agree with the small shifts observed when a 100 K cryogenic structure (blue bonds) is superimposed on the 298 K XFEL structure (gold bonds). (b) The populations of C67 alternate conformations are altered by the change in temperature between the synchrotron and XFEL datasets, as shown by difference electron density peaks near the sulfur atom (yellow). (c) The populations of alternate conformations of C101 are altered by change in temperature, as shown by peaks near the sulfur atom (yellow).

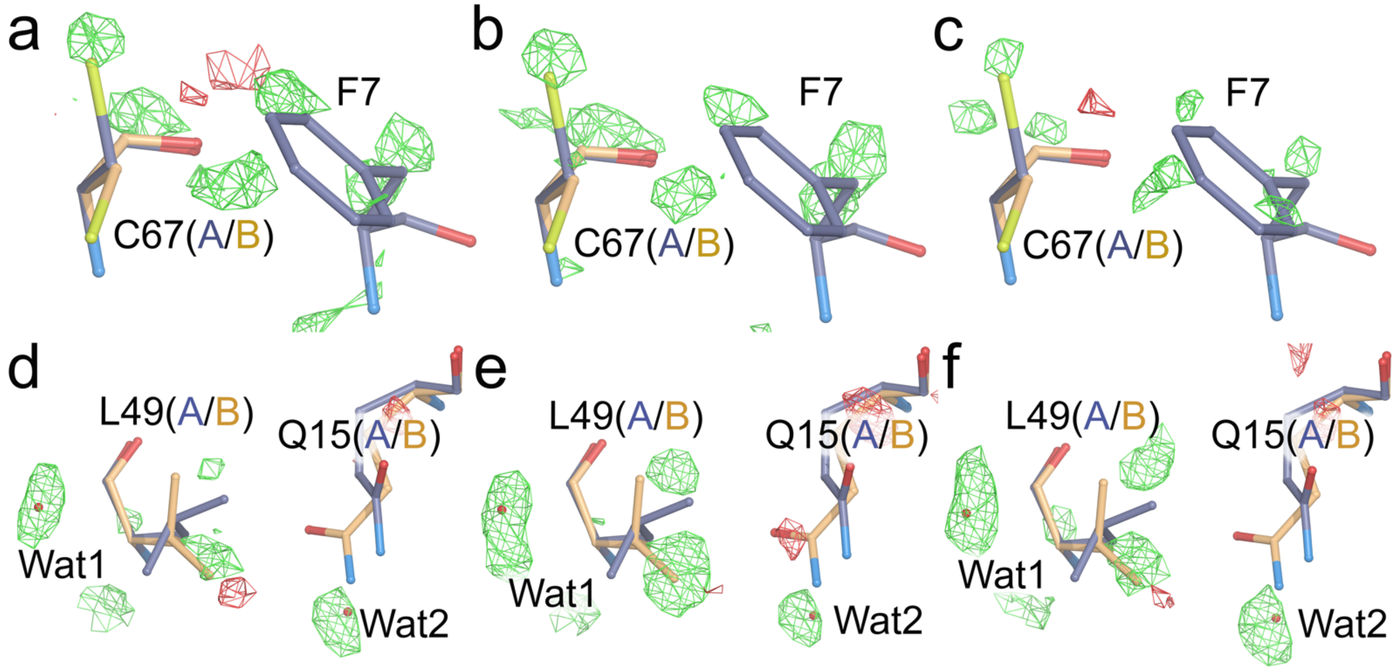


**Figure S3. Difference electron density features between synchrotron and XFEL datasets are reproducible.** $F_o$(274 K synchrotron)-$F_o$(298 K XFEL) isomorphous difference electron density maps are contoured at 2.8-3 $\sigma$ (corresponding to 0.17-0.18 $e^-$/Å$^3$) with positive features in green and negative ones in red. (a-c) show the same region near C67 with $F_o$(274 K synchrotron)-$F_o$(298 K XFEL) isomorphous difference electron density maps calculated from three independent synchrotron datasets. (d-f) show the region near L49 with $F_o$(274 K synchrotron)-$F_o$(298 K XFEL) isomorphous difference electron density maps calculated from three independent synchrotron datasets. The difference electron density features agree closely between the three datasets.

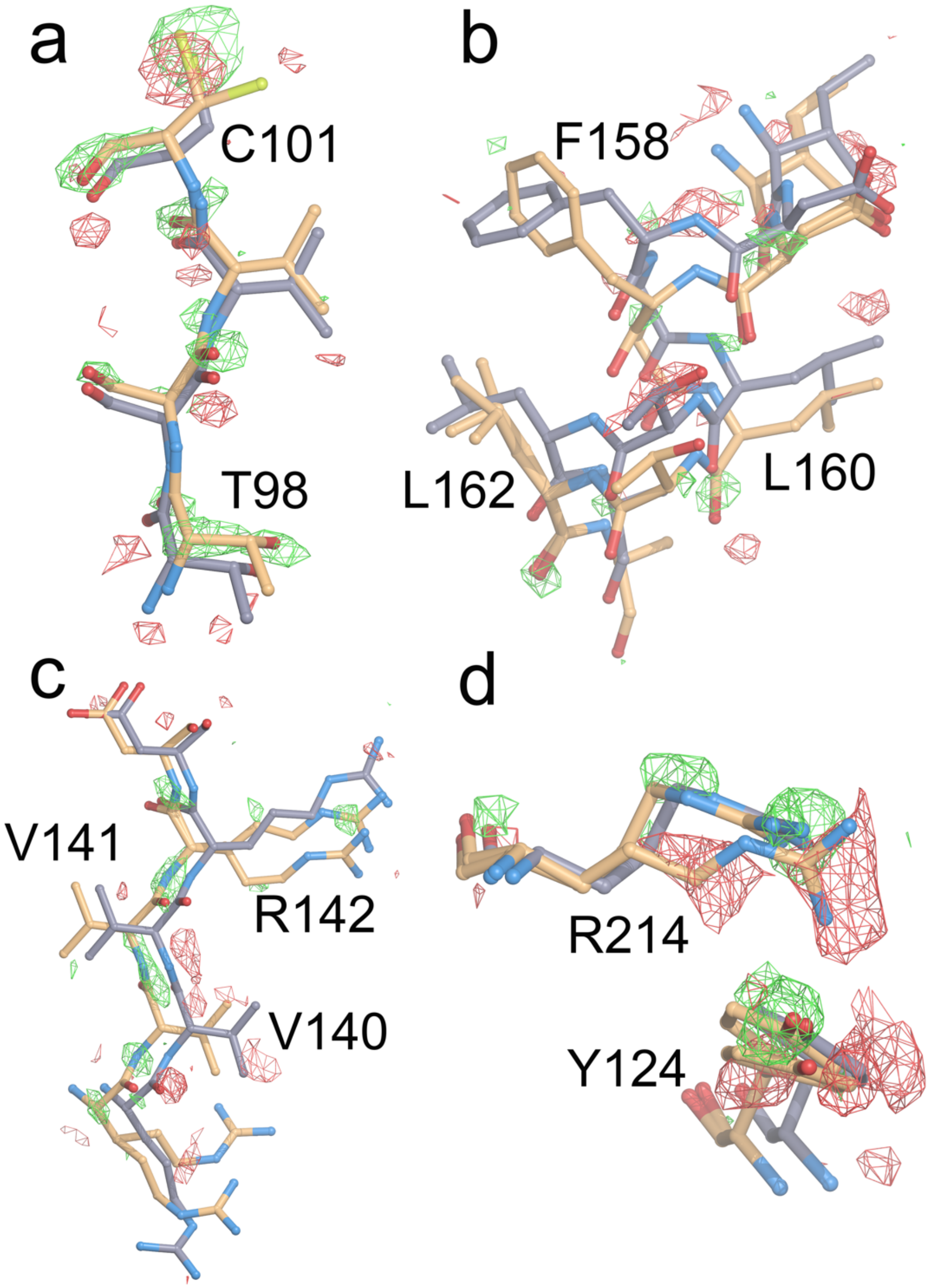


**Figure S4. The G150T mutation causes ICH to populate conformations that the wild-type enzyme samples during catalysis.** $F_o(15s)$-$F_o(0s)$ isomorphous difference electron density maps calculated for wild-type ICH from a prior experiment [1] are contoured at 2.7$\sigma$ (corresponding to 0.15 $e^-/Å^3$). Wild-type ICH is shown in slate gray and G150T ICH is shown in gold bonds. (a-d) Selected examples where the conformational shifts evident between the wild-type and G150T ICH structures agree well with the difference electron density map peaks observed during catalysis in wild-type ICH. In all panels, the G150T ICH structure (gold) overlaps with positive difference electron density peaks.

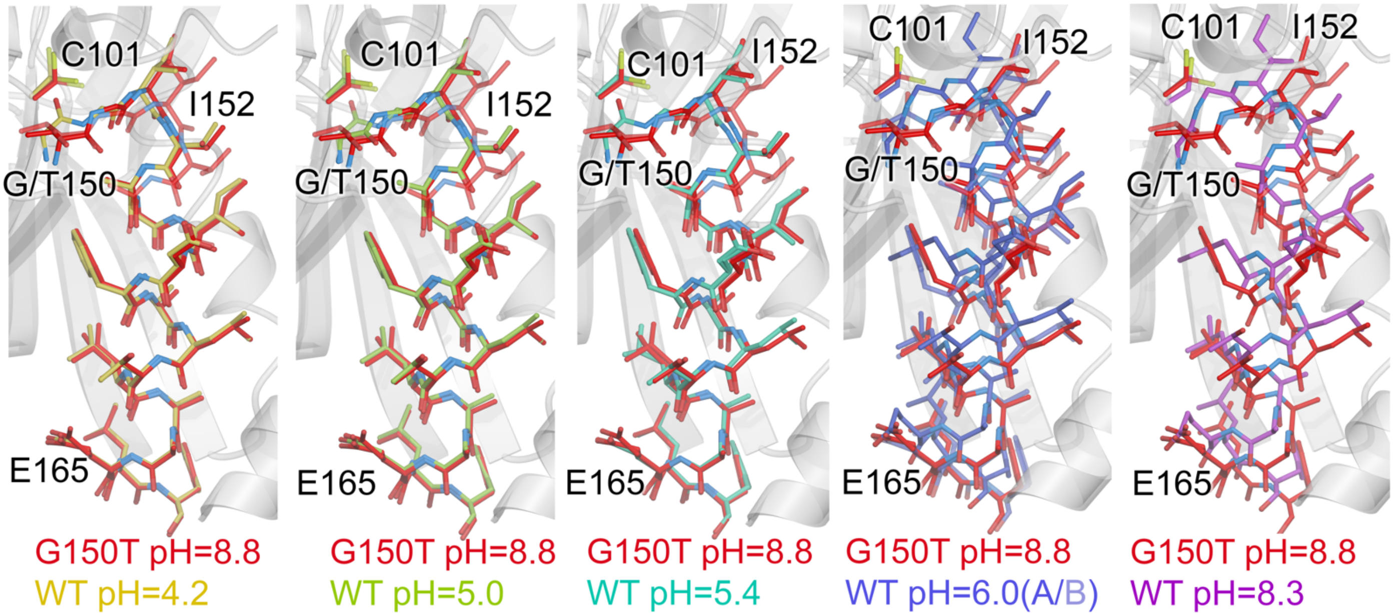


**Figure S5. pH-dependent shifts of the helix H conformation in wild-type ICH recapitulate changes resulting from the G150T mutation.** The structure of G150T ICH is shown in red bonds and structures of wild-type (WT) ICH at the indicated pH values are shown in bonds ranging from yellow (pH=4.2) to purple (pH=8.3). As pH decreases, wild-type ICH adopts a conformation that is highly similar to G150T ICH, consistent with changes expected from protonation of C101.

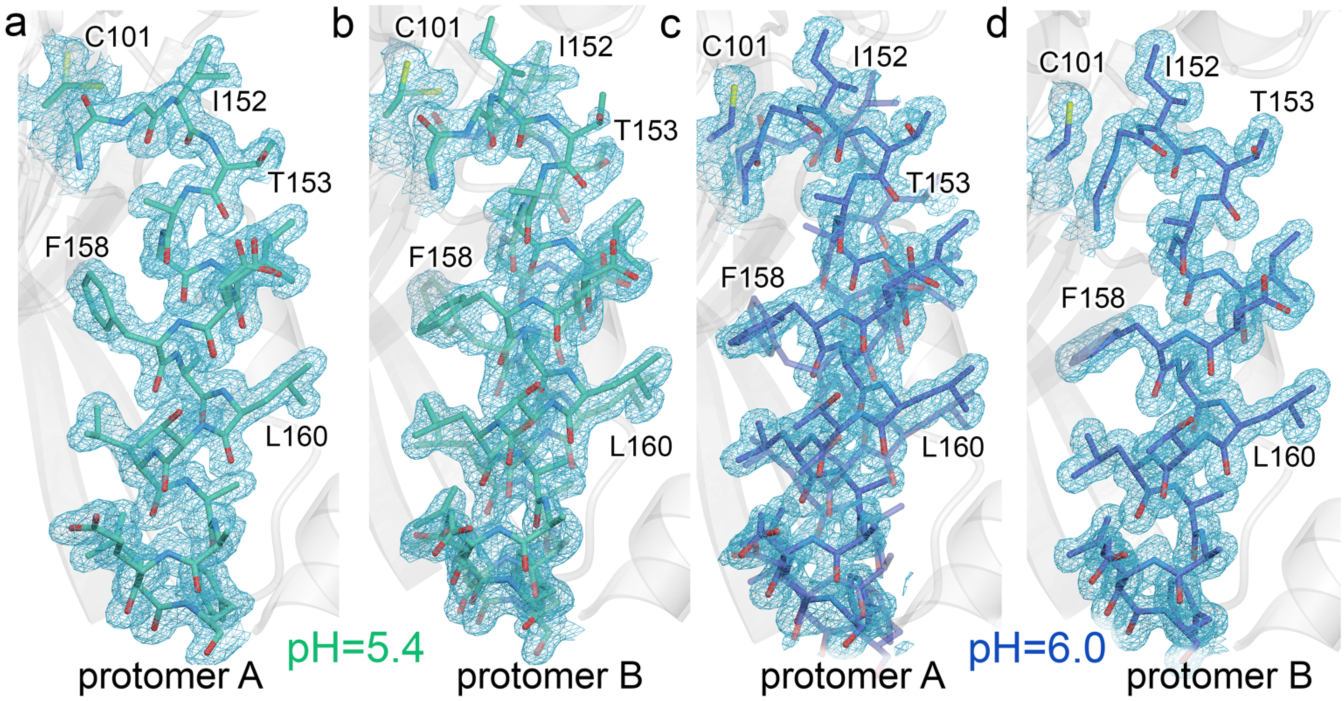


**Figure S6. pH-dependent conformational heterogeneity in helix H is asymmetric in the wild-type ICH dimer.** $2mF_o$-$DF_c$ electron density is shown at $0.9\sigma$ for wild-type ICH at pH=5.4 (green; a, b) and pH=6.0 (blue; c, d). The dominant conformer is shown in solid bonds and the minor conformer is semi-transparent. (a,b) At pH=5.4, protomer A is in the fully shifted conformation and protomer B is a mixture of shifted and strained helix conformations. (c,d) At pH=6.0, protomer A is a mixture of shifted and strained helix conformations and protomer B is in the fully strained conformation. In general, the helix in protomer A has a stronger tendency to sample the shifted (G150T-like) conformation.

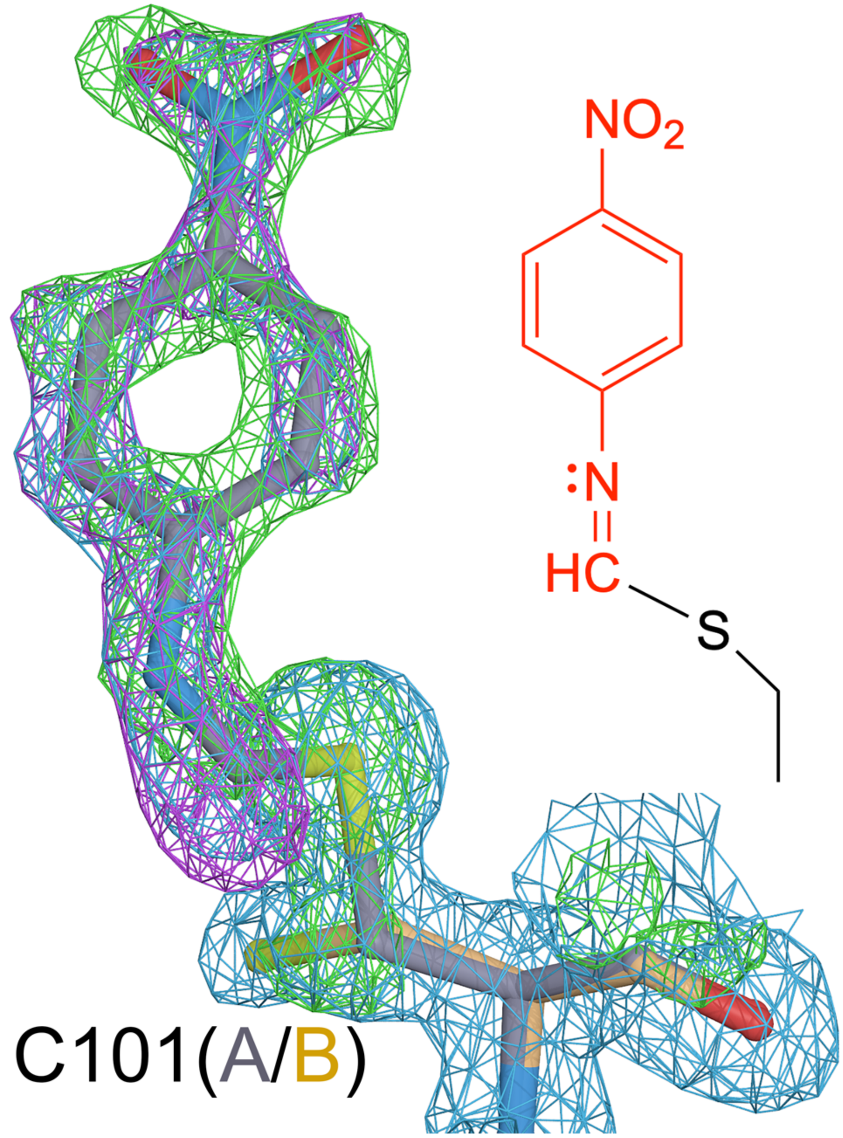


**Figure S7. Thioimidate intermediate forms in G150T ICH 30 seconds after mixing with p-NPIC substrate.** The $F_o(30s)$-$F_o(0s)$ isomorphous difference electron density map ($2.7\sigma$, purple), $mF_o$-$DF_c$ omit difference electron density map calculated before the model of the thioimidate was included ($3.0\sigma$, green), and the $2mF_o$-$DF_c$ omit electron density map calculated before the model of the thioimidate was included ($0.8\sigma$, blue) all show clear evidence for the covalent modification of C101. The chemical structure of the intermediate is shown to the right.

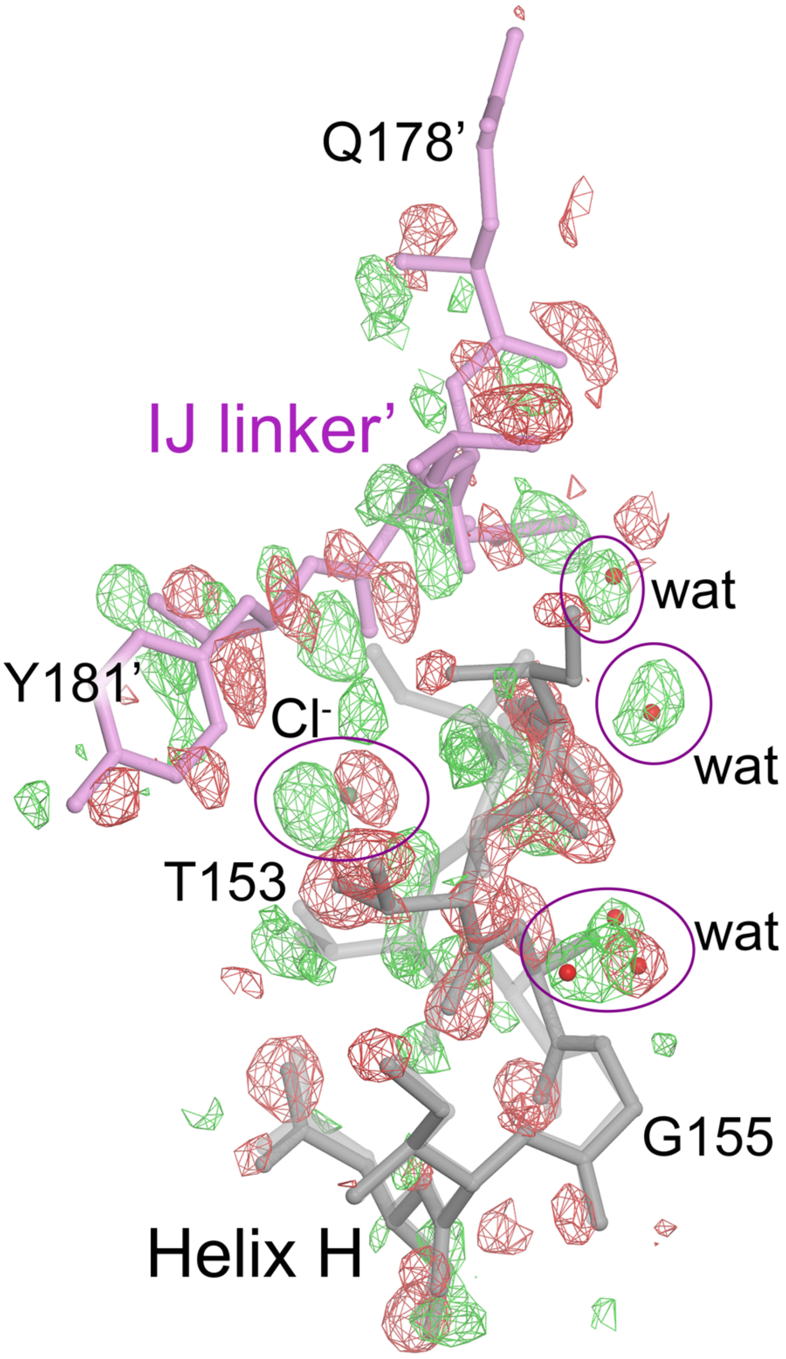


**Figure S8. Correlated changes in protein and solvent during catalysis in G150T ICH.** $F_o$(30s)-$F_o$(0s) isomorphous difference electron density map is contoured at 3.0$\sigma$ with positive features in green and negative ones in red. Helix H, the IJ linker region from the other protomer (purple, indicated with apostrophe), and surrounding solvent (red spheres, labeled “wat”) all display spatially correlated changes in conformation upon intermediate formation during catalysis.

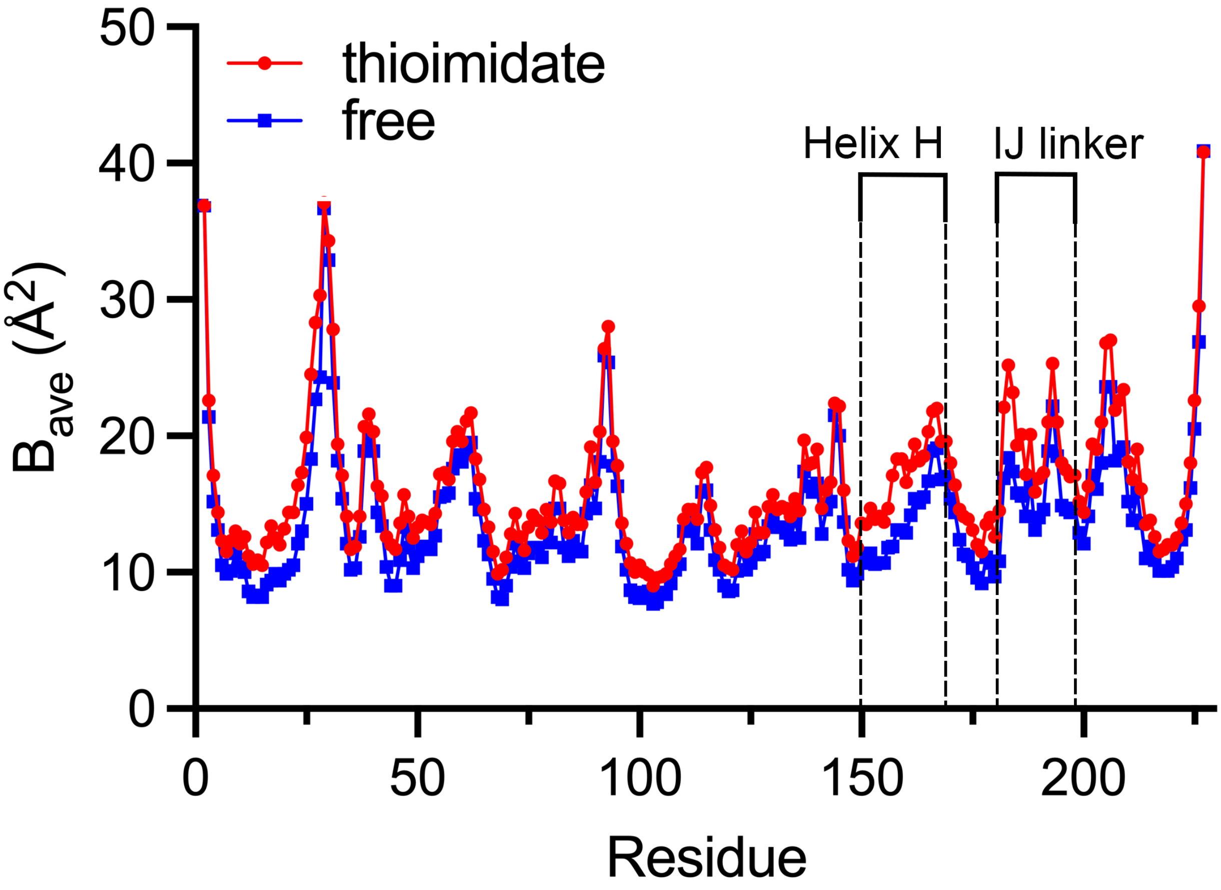


**Figure S9. Regions of elevated average ADPs in the G150T thioimidate intermediate.** Residue-averaged main chain values for the ADPs ($B_{ave}$) are plotted vs. residue number. Helix H and the IJ linker (labeled) have markedly higher average mainchain ADPs upon thioimidate formation during G150T ICH catalysis compared to the resting free enzyme.

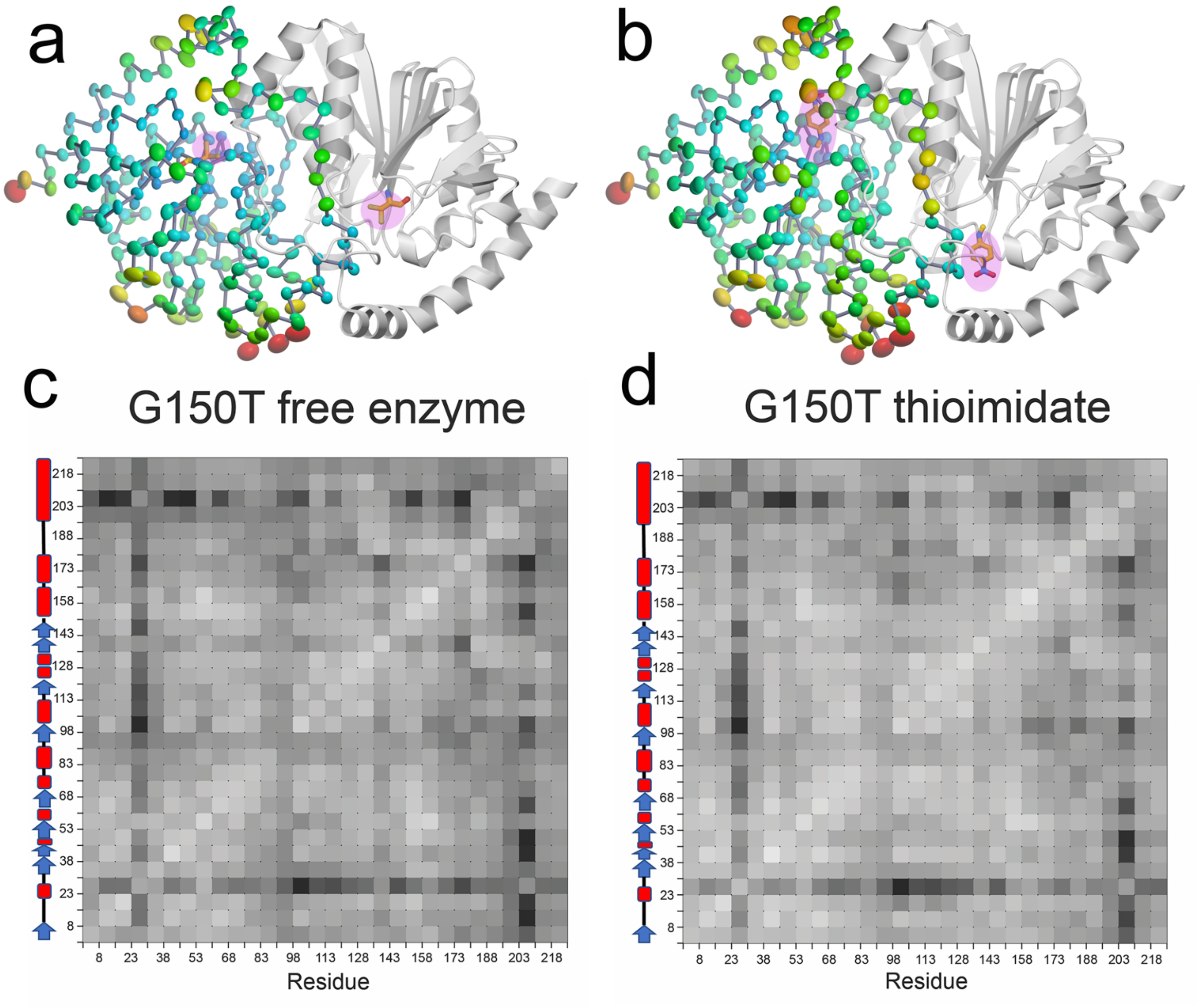


**Figure S10. Anisotropic ADPs in G150T ICH change in magnitude but not direction upon thioimidate formation.** (a,b) Anisotropic ADP ellipsoids for C$\alpha$ atoms at the 85% probability level are colored by magnitude, from blue (5 Å$^2$) to red (30 Å$^2$). Formation of the thioimidate intermediate (labeled; pink highlight) during catalysis elevates ADP magnitudes in the active site, the IJ linker, and helix H. Larger ADP magnitudes are indicated with larger ellipsoid volumes and warmer colors. (c,d) Rosenfield difference analysis [2] indicates no major changes in ADP directional preferences upon intermediate formation. Secondary structural elements in ICH are shown on the left to the Rosenfield matrices, with helices as red rectangles and strands as blue arrows. Rosenfield analysis determines the difference between the projections of two atoms' ADP ellipsoids onto the line joining them. Rigid body-like correlated motion of a pair of atoms would result in low difference values (lighter colors). The Rosenfield matrix is computed by averaging mainchain difference ADP projection values in 30 bins (~7 residues per bin).

**TABLE S1.** Crystallographic data statistics for synchrotron datasets

| Sample | G150T 100 K | G150T-1 274K | G150T-2 274K | G150T-3 274K | WT pH 4.2 | WT pH 5.0 | WT pH 5.4 | WT pH 6.0 | WT pH 8.3 |
|---|---|---|---|---|---|---|---|---|---|
| Diffraction source | SSRL 12-2 | SSRL 12-2 | SSRL 12-2 | SSRL 12-2 | SSRL 9-2 | SSRL 9-2 | SSRL 9-2 | SSRL 9-2 | SSRL 9-2 |
| Wavelength (Å) | 0.775 | 0.775 | 0.775 | 0.775 | 0.886 | 0.886 | 0.886 | 0.886 | 0.886 |
| Temperature (K) | 100 | 274 | 274 | 274 | 100 | 100 | 100 | 100 | 100 |
| Detector | Pilatus 6M | Pilatus 6M | Pilatus 6M | Pilatus 6M | Pilatus 6M | Pilatus 6M | Pilatus 6M | Pilatus 6M | Pilatus 6M |
| Space group | $P2_1$ | C2 | C2 | C2 | $P2_1$ | $P2_1$ | $P2_1$ | $P2_1$ | $P2_1$ |
| a, b, c (Å) | 55.56, 58.65, 68.87 | 72.19, 59.72, 56.30 | 72.20, 59.73, 56.37 | 72.13, 59.66, 56.17 | 57.04, 40.60, 83.28 | 57.05, 40.72, 83.43 | 56.53, 56.79, 68.77 | 56.75, 56.79, 68.41 | 56.87, 56.83, 68.24 |
| α, β, γ (°) | 90.00, 110.92, 90.00 | 90.00, 115.89, 90.00 | 90.00, 115.88, 90.00 | 90.00, 115.86, 90.00 | 90.00, 104.28, 90.00 | 90.00, 104.42, 90.00 | 90.00, 112.44, 90.00 | 90.00, 112.54, 90.00 | 90.00, 112.54, 90.00 |
| Mosaicity (°) | 0.08 | 0.10 | 0.14 | 0.08 | 0.12 | 0.13 | 0.15 | 0.10 | 0.06 |
| Resolution range (Å) | 38.87-1.00 (1.02-1.00) | 39.47-1.15 (1.17-1.15) | 35.21-1.20 (1.22-1.20) | 35.16-1.10 (1.12-1.10) | 40.35-1.50 (1.53-1.50) | 37.34-1.45 (1.47-1.45) | 38.45-1.33 (1.36-1.33) | 38.52-1.20 (1.22-1.20) | 38.57-1.02 (1.04-1.02) |
| Total no. of observations | 1035459 (45149) | 280457 (12317) | 254082 (11584) | 302439 (13882) | 323496 (14785) | 360375 (16343) | 547538 (28370) | 725100 (36449) | 1238270 (51309) |
| No. of unique observations | 218349 (10505) | 74206 (3528) | 66132 (3293) | 82868 (3983) | 58834 (2890) | 65010 (3157) | 89288 (4619) | 123914 (6174) | 201545 (10211) |
| Completeness (%) | 98.1(95.6) | 97.4 (94.3) | 98.3 (95.8) | 95.7 (93.4) | 98.9 (99.2) | 98.4 (97.5) | 97.3 (95.8) | 99.2 (97.5) | 99.4 (98.5) |
| Multiplicity | 4.7 (4.3) | 3.8 (3.5) | 3.8 (3.5) | 3.6 (3.5) | 5.5 (5.1) | 5.5 (5.2) | 6.1 (6.1) | 5.9 (5.9) | 6.1 (5.0) |
| ⟨I/σ(I)⟩ | 9.3 (0.5) | 12.0 (0.9) | 7.3 (1.1) | 11.8 (1.0) | 9.5 (0.9) | 10.1 (1.0) | 13.3 (0.8) | 9.8 (1.0) | 12.5 (0.9) |
| $CC_{1/2}$[1] | 0.999 (0.262) | 0.999 (0.366) | 0.997 (0.334) | 0.998 (0.334) | 0.999 (0.404) | 0.999 (0.427) | 0.999 (0.325) | 0.999 (0.435) | 0.999 (0.350) |
| $R_{meas}$ | 0.063 (2.850) | 0.052 (1.720) | 0.078 (2.243) | 0.057 (1.839) | 0.107 (2.150) | 0.097 (1.822) | 0.067 (2.504) | 0.096 (1.921) | 0.071 (1.944) |

[1] $CC_{1/2}$ [3] was used to determine the high-resolution cutoff.

**TABLE S2.** Crystallographic data statistics for XFEL datasets

| Sample | G150T apo XFEL | G150T 30s XFEL |
|---|---|---|
| Diffraction source | LCLS MFX | LCLS MFX |
| Wavelength (Å) | 1.033 | 1.033 |
| Temperature (K) | 298 | 298 |
| Detector | Rayonix MX340-XFEL | Rayonix MX340-XFEL |
| Space group | C2 | C2 |
| a, b, c (Å) | 72.13, 59.85, 56.14 | 72.14, 59.76, 56.11 |
| α, β, γ (°) | 90.00, 115.89, 90.00 | 90.00, 115.88, 90.00 |
| Resolution range (Å) | 22.00-1.30 (1.32-1.30) | 21.98-1.30 (1.32-1.30) |
| Total number of indexed images | 23990 | 22042 |
| Total no. of observations | 3133191 (43462) | 2240519 (42908) |
| No. of unique observations | 52803 (2583) | 52683 (2580) |
| Completeness (%) | 99.97 (100.0) | 99.97 (99.96) |
| Multiplicity | 59.33 (16.83) | 42.52 (16.62) |
| ⟨I/σ(I)⟩ | 3.49 (0.92) | 3.21 (0.96) |
| $CC_{1/2}$ | 0.964 (0.650) | 0.948 (0.666) |
| $R_{split}$[1] | 0.181 (0.494) | 0.189 (0.550) |

$R_{split}$ calculated as in [4].

### **TABLE S3.** Crystallographic refinement statistics

| Sample | G150T XFEL Free | G150T XFEL 30s | G150T cryo | G150T-1 RT | G150T-2 RT | G150T-3 RT | WT ICH pH 4.2 | WT ICH pH 5.0 | WT ICH pH 5.4 | WT ICH pH 6.0 | WT ICH pH 8.3 |
|---|---|---|---|---|---|---|---|---|---|---|---|
| PDB code | 8TSF | 8TSN | 8TSX | 8TSU | 8TSY | 8TSZ | 8TT0 | 8TT1 | 8TT2 | 8TT4 | 8TT5 |
| Temperature (K) | 298 | 298 | 100 | 274 | 274 | 274 | 100 | 100 | 100 | 100 | 100 |
| Refinement program | PHENIX 1.19.2-4158 | PHENIX 1.19.2-4158 | PHENIX 1.19.2-4158 | PHENIX 1.19.2-4158 | PHENIX 1.19.2-4158 | PHENIX 1.19.2-4158 | PHENIX 1.19.2-4158 | PHENIX 1.19.2-4158 | PHENIX 1.19.2-4158 | PHENIX 1.19.2-4158 | PHENIX 1.19.2-4158 |
| Resolution range (Å) | 22-1.30 (1.32-1.30) | 21.98-1.30 (1.32-1.30) | 38.87-1.00 (1.01-1.00) | 35.25-1.15 (1.17-1.15) | 35.21-1.20 (1.22-1.20) | 35.16- 1.10 (1.12-1.10) | 40.35-1.50 (1.52-1.50) | 32.78-1.45 (1.47-1.45) | 38.45-1.33 (1.35-1.33) | 38.52-1.20 (1.22-1.20) | 38.01-1.02 (1.03-1.02) |
| Completeness (%) | 99.98 (100) | 99.98 (100) | 95.92 (59) | 96.82 (88) | 98.13 (95) | 95.42 (93) | 98.62 (98) | 98.07 (96) | 96.98 (94) | 99.19 (97) | 98.69 (82) |
| No. of reflections | 52794 (2606) | 52673 (2590) | 213316 (4309) | 73920 (2347) | 66082 (2474) | 82841 (2642) | 58739 (2606) | 64920 (2569) | 89186 (2734) | 123860 (3786) | 200123 (5317) |
| No. of reflections, test set | 2600 (134) | 2591 (129) | 5143 (101) | 3801 (114) | 3331 (132) | 4122 (144) | 2905 (121) | 3233 (152) | 4336 (138) | 6020 (214) | 9758 (290) |
| $R_{work}$ | 0.1428 (0.2456) | 0.1546 (0.2802) | 0.1417 (0.3557) | 0.1196 (0.3113) | 0.1175 (0.2914) | 0.1161 (0.2732) | 0.1684 (0.3211) | 0.1556 (0.3060) | 0.1457 (0.3765) | 0.1288 (0.2911) | 0.1216 (0.2939) |
| $R_{free}$ | 0.1709 (0.2721) | 0.1835 (0.3109 ) | 0.1679 (0.4055) | 0.1405 (0.2862) | 0.1371 (0.3104) | 0.1338 (0.2629) | 0.1960 (0.3421) | 0.1818 (0.3441) | 0.1763 (0.3838) | 0.1581 (0.2891) | 0.1377 (0.3035) |
| No. of non-H atoms | | | | | | | | | | | |
| Protein | 1994 | 1963 | 4116 | 2002 | 2002 | 2002 | 3642 | 3567 | 4280 | 4518 | 4180 |
| Water | 163 | 155 | 447 | 178 | 178 | 181 | 407 | 373 | 433 | 537 | 538 |
| Heteroatom | 1 | 12 | 5 | 1 | 1 | 1 | 0 | 0 | 16 | 28 | 28 |
| Total | 2158 | 2130 | 4568 | 2181 | 2181 | 2184 | 4049 | 3940 | 4729 | 5083 | 4746 |
| Average R.M.S. deviations | | | | | | | | | | | |
| Bonds (Å) | 0.005 | 0.007 | 0.007 | 0.007 | 0.010 | 0.008 | 0.007 | 0.005 | 0.005 | 0.005 | 0.008 |
| Angles (°) | 0.827 | 0.986 | 0.965 | 0.880 | 1.029 | 0.972 | 0.771 | 0.746 | 0.856 | 0.890 | 1.020 |
| Average B factors ($\langle B_{iso} \rangle$;Å$^2$) | | | | | | | | | | | |
| Protein | 16.27 | 18.55 | 15.34 | 17.90 | 18.24 | 16.57 | 22.40 | 21.09 | 20.89 | 14.83 | 12.67 |
| Water | 35.80 | 36.68 | 32.98 | 37.05 | 36.83 | 33.67 | 30.43 | 34.29 | 37.08 | 32.15 | 27.33 |
| Average ADP anisotropy[1] | | | | | | | | | | | |
| Protein | 0.490 | 0.519 | 0.442 | 0.430 | 0.487 | 0.494 | 0.561* | 0.551* | 0.396 | 0.403 | 0.458 |
| Water | 0.436 | 0.452 | 0.394 | 0.357 | 0.413 | 0.404 | 1.000 | 0.390* | 0.394 | 0.366 | 0.394 |
| MolProbity clashscore[2] | 2.23 | 1.25 | 3.95 | 1.97 | 1.72 | 1.48 | 1.76 | 1.11 | 4.49 | 2.72 | 1.76 |
| Ramachandran plot | | | | | | | | | | | |
| Outliers (%) | 0.00 | 0.00 | 0.00 | 0.00 | 0.00 | 0.00 | 0.00 | 0.00 | 0.00 | 0.44 | 0.44 |
| Allowed (%) | 0.89 | 0.89 | 0.89 | 0.89 | 0.89 | 0.45 | 0.89 | 0.88 | 1.33 | 1.11 | 1.11 |
| Favored (%) | 99.11 | 99.11 | 99.11 | 99.11 | 99.11 | 99.55 | 99.11 | 99.12 | 98.67 | 98.45 | 98.45 |

[1]Anisotropy is defined as the ratio of the smallest to largest eigenvalue of the ADP tensor and was calculated using PARVATI [5]

[2]Defined as the number of close contacts (atom overlap > 0.4 Å) per 1000 atoms [6].
